# Deciphering the impact of cancer cell’s secretome and its derived-peptide VGF on breast cancer brain metastasis

**DOI:** 10.1101/2024.02.22.581537

**Authors:** Rita Carvalho, Liliana Santos, Inês Conde, Ricardo Leitão, Hugo R. S. Ferreira, Célia Gomes, Ana Paula Silva, Fernando Schmitt, Carina Carvalho-Maia, João Lobo, Carmen Jerónimo, Joana Paredes, Ana Sofia Ribeiro

**Author notes:** Ana Sofia Ribeiro **Adress: R. Alfredo Allen 208, 4200-135 Porto To whom correspondence may be addressed. Email:**. **Conflict of interest statement** The authors have no conflicting interests to disclose.

## Abstract

Brain metastases (BM) are one of the most serious clinical problems in breast cancer (BC) progression, associated with lower survival rates and a lack of effective therapies. Thus, to dissect the early stages of the brain metastatic process, we have searched for a brain-tropic metastatic signature on BC cells’ secretome, as a promising source for the discovery of new biomarkers involved in brain metastatic progression.

Therefore, six specifically deregulated peptides were found to be enriched in the secretome of brain organotropic BC cells. Importantly, these secretomes caused significant blood-brain barrier (BBB) disruption, as well as microglial activation, *in vitro* and *in vivo*. We identified the VGF nerve growth factor inducible as a brain-specific peptide, promoting BBB dysfunction similar to the secretome of brain organotropic BC cells. Concerning microglial activation, a slight increase was also observed upon VGF treatment.

In a series of human breast tumors, VGF was found to be expressed in both cancer cells and in the adjacent stroma. VGF-positive tumors showed a significant worse prognosis and were associated with HER2 overexpression and triple-negative molecular signatures. Finally, in a cohort including primary breast tumors and their corresponding metastatic locations to the lung, bone, and brain, we found that VGF significantly correlates with the brain metastatic site.

In conclusion, we found a specific BC brain metastatic signature, where VGF was identified as a key mediator in this process. Importantly, its expression was associated with poor prognosis for BC patients, probably due to its associated increased risk of developing BM.

## Introduction

Breast cancer (BC) is the most common malignancy and leading cause of cancer death in women due to distant metastases to the bone, lung, liver, and brain (1). According to Paget’s theory, metastases are a non-random process and are instead determined by the interaction between cancer cells (the seeds) and the metastatic microenvironment (the soil) (2–4). For example, BC expresses chemokine receptors, namely CXCR4 and CCR7, which pair with chemokine ligands expressed in lymph nodes (CXCL12) and lung (CCL21), thus determining the destination of metastases (5). Several studies already showed that cancer cells modulate metastatic niches way before they reach the target organ. Several efforts are being made to uncover and characterize the molecular factors that are secreted by cancer cells, aiming to identify metastatic biomarkers. Long before intravasation, cancer cells communicate with distant metastatic organs, mainly by secreting soluble factors and/or extracellular vesicles (EVs), that recruit immune cells, promote resident cells activation, and extracellular matrix (ECM) remodeling, thereby preparing a supportive and receptive pre-metastatic niche (PMN) to the growth and survival of cancer cells (6–9). These molecular factors released by cancer cells collectively compose the “cancer cell secretome”, which is a promising source of molecular targets involved in the crosstalk between the primary tumor and the future metastatic site (8, 10). Importantly, PMNs may influence the success of metastatic colonization, and understanding its formation and regulation may help to improve the development of effective therapies.

Although BC survival rates have improved significantly in recent years, owing to advancements in surveillance and more effective systemic treatments, 10% to 30% of patients with metastatic BC will develop brain metastases (BM) during the course of their disease (11–14). Breast cancer brain metastasis (BCBM) are a major cause of poor quality of life and the prognosis of patients remains dismal, with survival typically measured in months (15–17). Current treatments of BM include surgery, stereotactic radiotherapy, whole brain radiotherapy and systemic therapy. However, no effective chemotherapeutic options for BM are available as opposed to treatments for lung and bone metastases. It is known that BC patients with HER2 overexpression and triple-negative molecular subtypes metastasize more frequently to the brain, suggesting that these cancer cells have specific molecular signatures that award their intrinsic advantage to survive in the brain tissue (11, 18). The unique composition of the brain, including the blood-brain barrier (BBB) and specialized resident cells, helps to create a highly protected environment, which is essential for its proper function and long-term health. Thus, cancer cell-brain metastatic niche interaction must present distinct properties compared with other metastatic organs. To prevent and treat BM, it is critical to identify new molecular drivers responsible for the communication between cancer cells and the brain microenvironment.

David Lyden’s research have already demonstrated that exosomes secreted by cancer cells contain specific molecules that allow them to home on target metastatic organs, a process known as organotropism (9, 19, 20). Particularly, in the brain, it was recently demonstrated that CEMIP exosomal protein pre-condition the brain microenvironment, enhancing cancer cell outgrowth (21). In fact, important efforts have been made to identify players involved in this process and to understand how cancer cells directly modulate the brain PMN. However, it is still an unexplored and very challenging area of research.

Therefore, in this work, we aimed to decipher the soluble factors secreted by BC cells responsible for the remodeling of the brain PMN. Our data demonstrated that the secretome of brain organotropic BC cells plays an important role in remodeling the brain PMN, namely by affecting the BBB integrity and microglia activation. Interestingly, we identified a specific brain secretome signature, being VGF (also known as VGF nerve growth factor inducible) one of the peptides specifically deregulated in the secretome of brain organotropic BC cells. Noteworthy, we validated its impact in the remodeling of the brain PMN and its potential to be a novel clinical predictive and prognostic biomarker for BM.

## Materials and Methods

### Cell Culture

The parental BC cell line MDA-MB-231 (parental 231) was obtained from ATCC (American Type Culture Collection, Manassas, VA, USA), while its organotropic BC variants lung (231.lung) (22), bone (231.Bone) (23), brain (231.Brain) (24) and brain with HER2 overexpression (231.Brain.HER2) (25) clones were obtained from J Massagué (MSKCC) and P. Steeg (NCI) **(Supplementary Figure S1)**. The human cerebral microvascular endothelial cell line (hCMEC/D3), was kindly provided by Pierre-Olivier Couraud (Institute Cochin, Université René Descartes, Paris, France). The human microglial clone 3 cell line (HMC3) was purchased from ATCC (ATCC^®^ CRL-3304™). Details are provided in **Supplementary Materials and Methods**.

### Secretomes Preparation

For *in vitro* assays, we plated BC cells (3x10^5^ cells in a 6 well-plate) embedded in collagen type ӏ (Merck) to mimic the breast ECM.

To study the secretomes impact *in vivo*, we plated BC cells (6x10^6^ cells) embedded in collagen type I in a T175 flask **(Supplementary Figure S2)**. Details in **Supplementary Materials and Methods**.

### In vitro evaluation of brain endothelial cell monolayer integrity

Cell monolayer integrity was determined by measuring the transendothelial flux of fluorescent 4kDa macromolecule across the endothelial cells (ECs) and the transendothelial electrical resistance (TEER) as previously described (26). Details about protocol and treatments are provided in **Supplementary Materials and Methods**.

### In vitro microglial phagocytosis assay

To evaluate phagocytic capacity, microglia cells were incubated with 0.0025% (w/w) 1 μm fluorescent latex beads for 75 minutes. After incubation, the total of microglia cells and the number of ingested beads per cell were counted using ImageJ (27). Details in **Supplementary Materials and Methods**.

### Mice secretome induced-model

Nude mice were pre-treated with secretomes (140 μg) or TLQP-21 (4.5 mg/kg; Sigma-Aldrich T1581) via intraperitoneal injection to assess their impact on BBB integrity and microglia modulation.

Mice follow-up was performed as previously described (28). All the experiments were conducted with the application of the 3Rs (replacement, reduction, and refinement) (JP_2016_02 Project, animal ethics committee, and animal welfare body of i3S) (29). Details are provided in **Supplementary Materials and Methods**.

### Immunofluorescence assay

Cells were fixed with 4% paraformaldehyde (PFA) for 20 minutes. Brain tissue was postfixed in 4% PFA for 24 hours. The protocol and antibodies can be found in the **Supplementary Materials and Methods**.

### Western Blot

Western blot was performed as described previously (30). Details about protocol and antibodies can be found in the **Supplementary Materials and Methods**.

### Proteomic analysis

100 μg of protein were processed for proteomics analysis following the solid-phase-enhanced sample-preparation (SP3) protocol as described by Hughes et al (31). Additional details are provided in **Supplementary Materials and Methods**.

### Kaplan Meier plotter survival analysis for VGF mRNA

Kaplan Meier (KM) plotter online survival analysis tool (https://kmplot.com) was used to assess the impact of VGF mRNA levels on overall survival (OS) in BC patients. Additional details are provided in **Supplementary Materials and Methods**.

### Primary Breast Cancer Series

A series of 218 primary breast carcinomas diagnosed between 1978–1992, were retrieved from the Pathology Department, Hospital Xeral-Cíes, Vigo, Spain. Patient follow-up information was available for the 218 **(Supplementary Table S1)**, with a maximum follow-up of 120 months after diagnosis. The tumors were characterized for clinical and pathological features, as previously described (32). All analyses were performed according to the reporting recommendations for tumor MARKer prognostic studies (REMARK) recommendations for prognostic and tumor marker studies. Details are provided in **Supplementary Materials and Methods.**

### Breast Cancer Metastases series

We used two complementary paired primary and metastatic BC series **(Supplementary Table S1).** The first one was retrospectively collected from the archives of the Department of Pathology of the Portuguese Oncology Institute of Porto (IPO Porto) and includes primary breast tumors that metastasized to the lung (n=17), bone (n=56), and brain (n=4). The second one is an exclusive BCBM series with paired primary breast tumors and their corresponding brain metastasis, which was retrospectively collected from Barretos Cancer Hospital, Brazil.

The present study was conducted with the approval of the Ethical Commission from both cancer centers, under the national regulative law for the usage of biological specimens from tumor banks, where the samples are exclusively available for research purposes in retrospective studies (Ethical approvals: Portuguese Oncology Institute of Porto (CES. 64/023) and Barretos Cancer Hospital/Fundação Pio XII (2-777-372). Details are provided in **Supplementary Materials and Methods.**

### Immunohistochemistry

Immunohistochemistry for IBA1 and VGF were performed in 3 µm sections. Further information regarding the protocol and antibodies can be found in the **Supplementary Materials and Methods**.

### Immunohistochemical evaluation

The expression of VGF was independently evaluated by one pathologist (F.S.) based on grading systems previously established for other markers (33, 34). Details are provided in **Supplementary Materials and Methods.**

### Statistical analyses

Prism software (GraphPad,v9.0) and IBM® SPSS® Statistics v.26 were used for statistical analysis, as detail in **Supplementary Materials and Methods**.

## Results

### The secretomes of brain organotropic breast cancer cells affect BBB

#### Integrity

Our goal was to evaluate the impact of the secretomes produced by brain organotropic BC cells in the preparation of the brain PMN, with a particular emphasis on BBB permeability and microglia activation. As models, brain organotropic BC cells (without or with HER2 overexpression), derived from the parental 231, were used **(supplementary material, Figure S1)**. To control for brain metastatic specificity, the results were always compared to data obtained with 231.Lung and 231.Bone organotropic BC cells.

To investigate the impact of the secretome of brain organotropic BC variants on the remodeling of the brain PMN, we started by evaluating BBB integrity. We found that the secretome of both brain organotropic BC variants show a specific and significant increase in cell permeability **(Figure 1A)** and decrease TEER **(Figure 1B)** of hCMEC/D3 monolayers. Interestingly enough, this significant effect was not seen with the secretome of the parental 231, nor with the ones from the non-brain organotropic BC variants. By immunofluorescence analyzes, we observed a significant decrease in β-catenin expression in hCMEC/D3 cells, in the presence of the secretomes of both brain organotropic BC variants **(Figure 1C-D)**, indicating an *in vitro* disruption of intercellular junctions.

**Figure 1.**
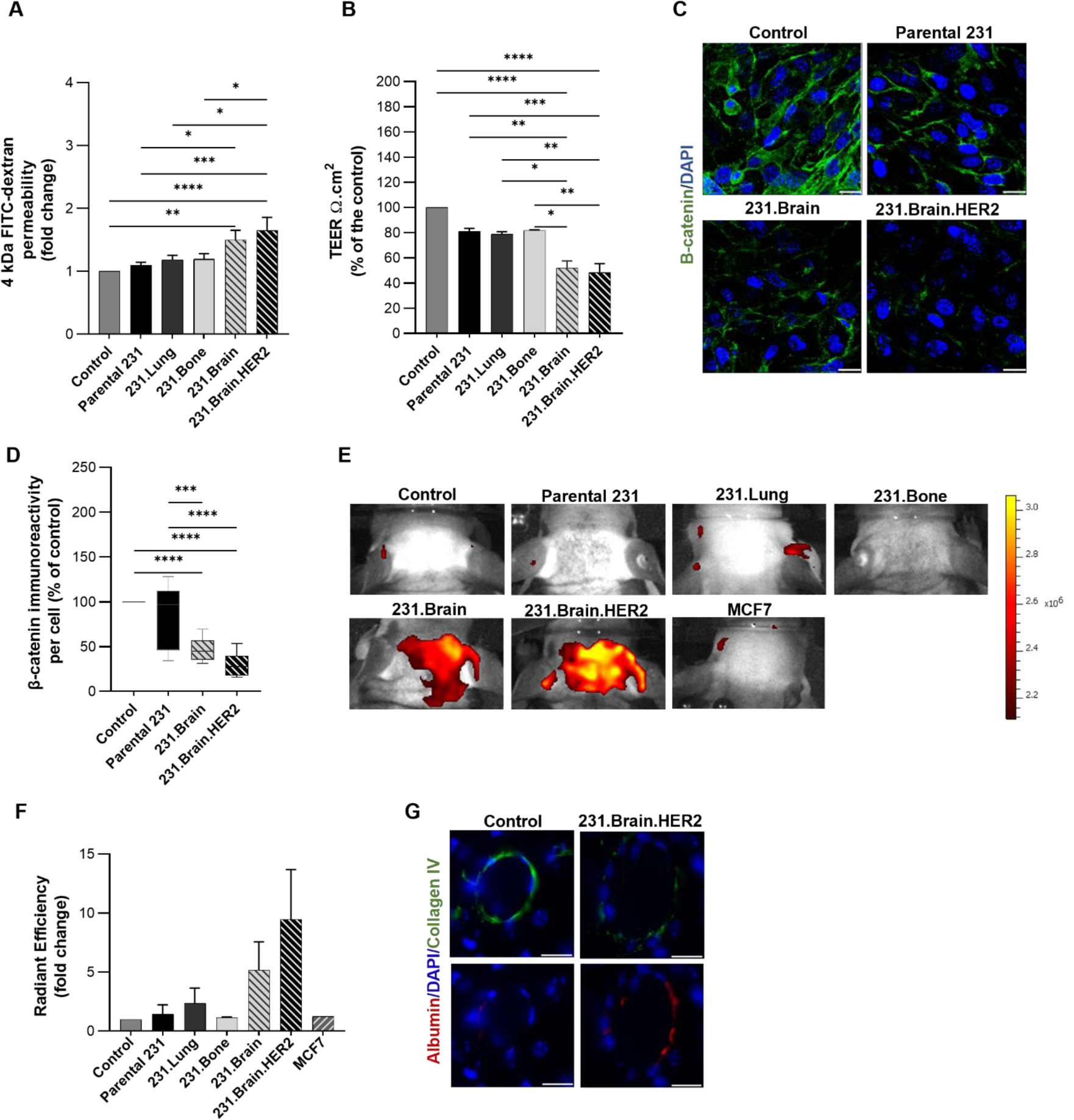
The secretome of brain organotropic breast cancer cells promotes BBB disruption. (A-D) Human cerebral microvascular endothelial cells (ECs) (hCMEC/D3 cells) monolayers were treated with serum-free medium (control condition) and secretomes collected from the parental 231 and their organotropic breast cancer variants. (A) *In vitro* permeability using a 4kDa FITC-dextran and (B) transendothelial electrical resistance (TEER) assay after 24 hours of treatment. (n=4 biological replicates; the results are expressed as mean ± SEM). (C) Representative confocal images of β-catenin. β-catenin: green, DAPI: Blue. (Scale bar corresponds to 20 μm). (D) Quantification of β-catenin. (n=3 biological replicates; the results are expressed as mean ± SEM). (E) *In vivo* near-infrared fluorescent brain imaging after 15 days of treating the mice with serum-free medium (control condition), the secretome of parental 231 and their organotropic breast cancer cells, as well as the secretome of MCF7 (a non-metastatic breast cancer cell line as a negative control). Representative images of mice brains acquired at 30 minutes post-injection of Cy7.5-dextran 5 kDa. (F) Quantification of radiant efficiency in equal-sized regions of interest (ROI) corresponding to the top of the skull (2/3 animals/group; the results are expressed as mean ± SEM). (G) Representative confocal images of collagen ІV (a marker of the basal membrane that gives support to brain vessels) and albumin (a marker of BBB disruption) staining after treatment with serum-free medium (control condition), as well as with the secretome of 231.Brain.HER2. Collagen ІV: green, albumin: red, DAPI: blue. (Scale bar corresponds to 20 μm). Statistical significance was assessed using one-way ANOVA.*P < 0.05, **P < 0.01, ***P < 0.001, ****P < 0.0001.

These results were further confirmed by the *in vivo* mice secretome-induced model. After priming mice with secretomes for 15 days, we showed a significant and specific impact of the secretome of both brain organotropic BC variants in BBB permeability, shown by an increased accumulation of a fluorescent dye in the brain, when compared with the secretome of the parental 231, MCF7 (non-metastatic BC cells) and serum-free DMEM (control condition) **(Figure 1E-F)**. Although the secretome from the non-brain organotropic BC variants also showed a small impact on the *in vivo* permeability after region of interest (ROI) measurement **(Figure 1F)**, this effect was more pronounced and specific when nude mice were pre-treated with the secretome from both brain organotropic BC cells. Concomitantly, the brain from these animals showed a decrease in collagen IV and an increase in albumin expression in the pre-frontal cortex when pre-treated with the secretome of 231.Brain.HER2 overexpression **(Figure 1G)**, indicating structural and functional changes in the BBB integrity. Taken together, these data provide strong evidence that the secretome of brain organotropic BC cells cause BBB disruption, thus promoting the passage of molecular factors through this unique brain barrier.

#### The secretomes of brain organotropic breast cancer cells promote microglia activation

Some studies demonstrated that microglia can be recruited and activated during brain metastatic colonization (35–41). However, little is known about its role in the remodeling of the brain microenvironment in the early stages of the metastatic cascade, more specifically during the formation of the brain PMN. In order to understand the impact of the secretomes of brain organotropic BC cells on microglia activation, HMC3 were pre-treated with the secretomes collected from the parental 231 and their organotropic BC variants to evaluate their phagocytic capacity, as well as Stat3 phosphorylation.

Our *in vitro* data demonstrate that the secretome of both brain organotropic BC cells promote microglia activation since this treatment significantly increased both phagocytic capacity **(Figure 2A and supplementary material, Figure S3A)**, as well as Stat3 phosphorylation **(Figure 2B-C)** of microglia cells when compared with the control condition. Nevertheless, the secretome of the other non-brain organotropic variants also induced microglia activation, demonstrating that the previously observed specificity for the BBB is not occurring in this setting.

**Figure 2.**
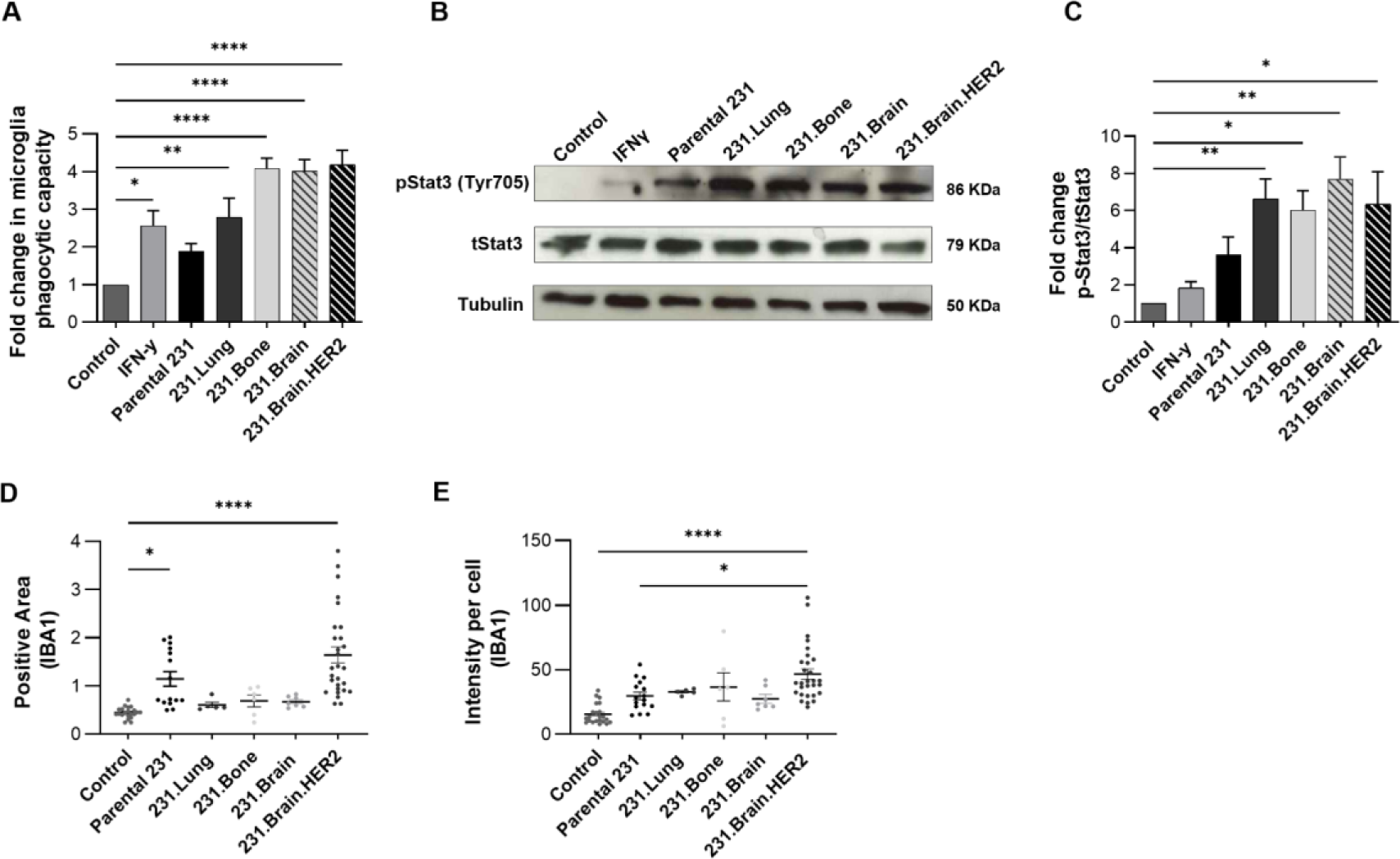
The secretome of brain organotropic breast cancer cells impacts microglia activation. (A) Quantification of microglia phagocytic capacity after treatment with serum-free DMEM (control condition), IFN-y (positive control), as well as with the secretomes of parental 231 and their organotropic breast cancer variants for 8 hours. (n=4 biological replicates; the results are expressed as mean ± SEM). (B) Western blot of Stat3 phosphorylation (pStat3 Tyr705) and total Stat3 (tStat3) in HMC3 cell lysates after 8 hours of treatment with the serum-free DMEM (control condition), IFN-y (positive control), as well as with the secretomes of parental 231 and their organotropic breast cancer variants. (C) Quantification of Stat3 phosphorylation over total Stat3 normalized to tubulin. (n=3 biological replicates; the results are expressed as mean ± SEM). (D) Quantification of microglia positive area and (E) staining intensity per microglia cell in the cerebral cortex of mice after 15 days of treatment with serum-free DMEM (control condition), as well as with the secretomes from the parental 231, and their organotropic breast cancer variants (IBA1-a marker of microglia cells). (2 mice/per group, 6-8 cerebral cortex independent areas; the results are expressed as mean ± SEM). Statistical significance was assessed using one-way ANOVA.*P < 0.05, **P < 0.01, ****P < 0.0001.

To validate these findings, we evaluated the role of the several secretomes on microglia cell population in the cerebral cortex from mouse models. Through immunohistochemical analysis for IBA1 (a microglia marker) **(supplementary material, Figure S3B)**, microglia cells were counted in the cerebral cortex, and the positive area and staining intensity by each cell was quantified, in order to measure their degree of activation. Interestingly, we demonstrated that the secretome of parental 231 significantly increased the positive area; however, the results were more evident with the secretome of 231.Brain.HER2 **(Figure 2D)**. In addition, this secretome significantly increased the staining intensity per cell **(Figure 2E)**, an effect not observed with the secretome of parental 231 and the other organotropic BC cells. These data suggest that factors present in the secretome of 231.Brain.HER2 trigger a more prominent response by microglial cells during the brain PMN formation.

#### The identification of a brain-specific protein secretome signature

In order to identify a specific brain secretome signature, high-throughput proteomic Label-Free quantitation analysis of the secretome of parental 231 and their organotropic BC variants was performed. We identified a total of 9064 peptides, but only 341 peptides were differentially expressed in the secretome of all organotropic BC cells compared with the secretome of the parental 231. In detail, within these peptides, 120 were differentially expressed in the secretome of 231.Lung, 75 in 231.Bone, 76 in 231.Brain and 70 in 231.Brain.HER2. We could observe a specific secretome signature for each metastatic location, as observed by unsupervised hierarchical clustering analysis **(supplementary material, Figure S4A)**. Interestingly, by principal component analysis (PCA), we observed that the secretome of parental 231 and the secretome of 231.Lung clusters close to each other, whether the secretome of both brain organotropic BC variants are more similar to the secretome of 231.Bone **(supplementary material, Figure S4B)**. To obtain the specific deregulated peptides present in the secretome of both brain organotropic BC variants, we intersected the lists of the significantly deregulated peptides. From this analysis, we identified 6 common peptides differentially expressed in both brain organotropic BC variants **(Figure 3A)**. Out of these, we focus on VGF due to its role on brain-related disorders. We further validated that VGF (full-length) was significantly enriched both at the cell level and in the secretome of both brain organotropic BC cells **(Fig. 3C-D and supplementary material, Figure S4 C)**.

**Figure 3.**
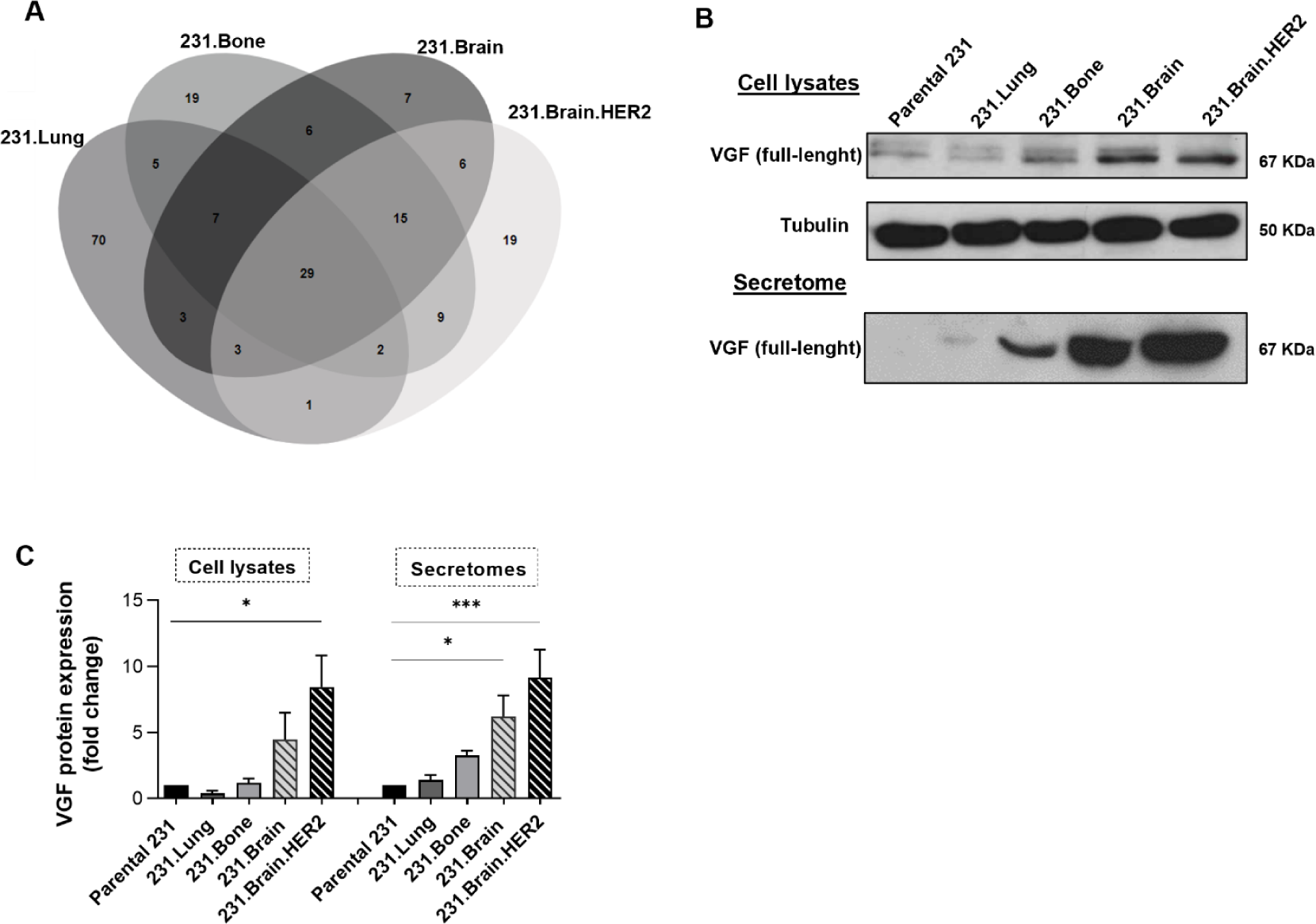
Brain organotropic breast cancer cells show a specific secretome signature. (A) Venny diagram to obtain the specific deregulated peptides in the secretome of both brain organotropic breast cancer variants. (B) Western Blot analysis to evaluate the expression and the presence of VGF in cell lysates and in the secretomes. (C) Western blot quantification. (n=4 and 8 biological replicates; the results are expressed as mean ± SEM). Statistical significance was assessed using one-way ANOVA.*P < 0.05, ***P < 0.001.

#### VGF promotes BBB disruption and microglia activation

We next investigated the role of VGF on the remodeling of the brain PMN, starting by its impact on BBB structure and function. We pre-treated hCMEC/D3 cell monolayer with TLQP-21, a bioactive C-terminal VGF-derived peptide for 24 hours. Interestingly, we found similar permeability and TEER values as observed with the secretome of 231.Brain.HER2 when compared with the control condition **(Figure 4A-B)**. Since TLQP-21 binds to the complement component 3a (C3a) receptor-1 (C3aR1), which is expressed in brain ECs (42), we further used a C3aR antagonist in the following experiments. Thus, we co-exposed hCMEC/D3 to the endogenous VGF secreted by 231.Brain.HER2 or to the exogenous VGF in combination with the selective antagonist for C3aR. Interestingly, we verified that the increase of endothelial permeability and the decrease of TEER values induced by exogenous and endogenous VGF were significantly prevented, proving that C3aR is mediating VGF-induced BBB hyperpermeability. Furthermore, we observed a significant decrease in β-catenin and ZO-1 **(Figure 4C-E)** protein levels in EC monolayers after treatment with the exogenous TLQP-21 peptide similar to what was observed with the secretome of 231.Brain.HER2. Accordingly, this result was also blocked in the presence of the C3aR antagonist.

**Figure 4.**
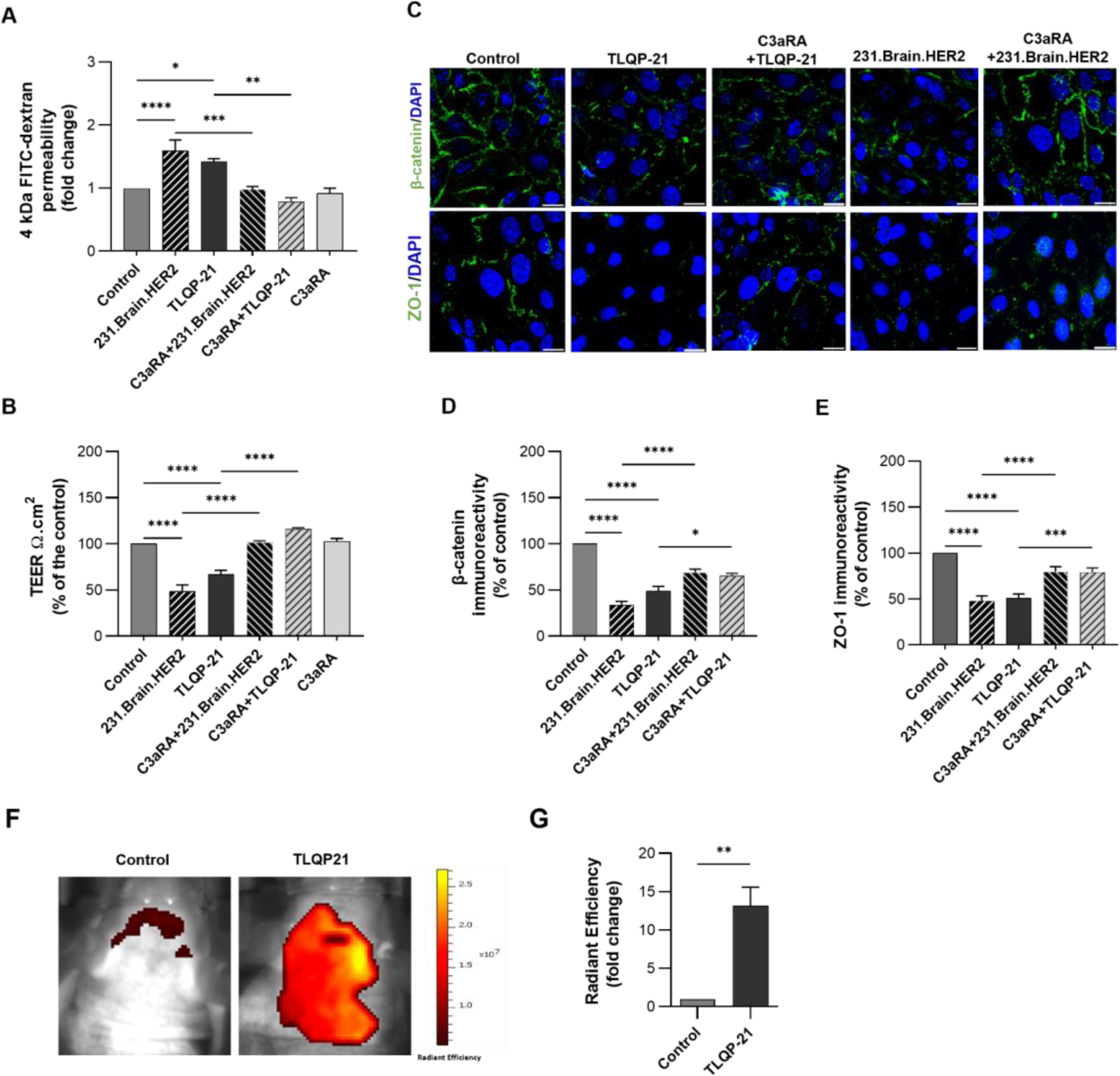
VGF promotes blood-brain barrier disruption. (A-B) Endothelial cells (ECs) monolayers were treated with TLQP-21 (100nM), a VGF derived-peptide, and with a secretome of 231.Brain.HER2 in the absence and presence of the C3a receptor antagonist (10µM) in order to evaluate (A) ECs monolayer permeability using a 4kDa FITC-dextran and (B) TEER after 24 hours of exposure. (n=4 biological replicates; the results are expressed as mean ± SEM). (C) Representative confocal images of β-catenin and ZO-1 immunoreactivity. β-catenin/ZO-1: green, DAPI: Blue. (Scale bar corresponds to 20 μm). (D-E) Quantification of β-catenin and ZO-1 expression. (F-G) *In vivo* dynamic near-infrared fluorescent imaging after 3 days of treatment with serum-free DMEM (control condition) and TLQP-21 (4.5 mg/kg). (F) Representative images of mice brains acquired at 30 minutes post-injection of Cy7.5-dextran 5 kDa. (G) Quantification of radiant efficiency in equal-sized ROI corresponding to the top of the skull. (4 animals/group; the results are expressed as mean ± SEM). Statistical significance was assessed using one-way ANOVA.*P < 0.05, **P < 0.01, ***, P < 0.001, ****P<0,0001

The same specific effect was also observed for microglia activation. Pre-treatment with TLQP-21 significantly increased the phagocytic capacity **(Figure 5A and supplementary material, Figure S5A)** and Stat3 phosphorylation **(Figure 5B-C)** of microglia cells, when compared with control condition. Interestingly, this effect was abolished when microglia cells were pre-treated with the C3aR antagonist, inhibiting VGF induced effects **(Figure 5A-C and supplementary material, Figure S5A)**.

**Figure 5.**
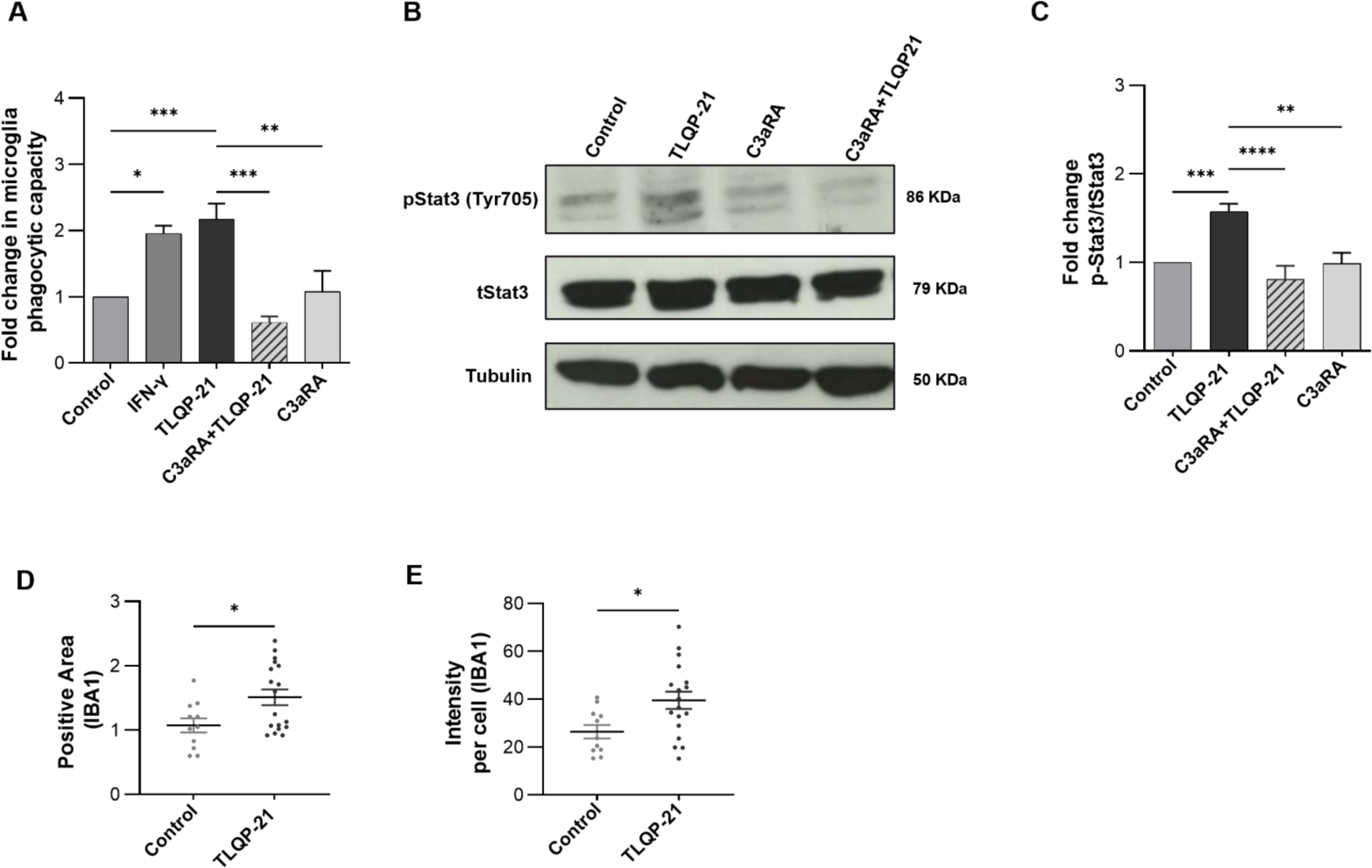
VGF impacts microglia activation. (A) Quantification of microglia phagocytic capacity after treatment with serum-free medium (control condition) and TLQP-21 (100nM) in the absence and presence of the C3a receptor antagonist (10µM) for 8 hours. (n=3 biological replicates; the results are expressed as mean ± SEM). (B) Western blot of Stat3 phosphorylation (pStat3 Tyr705) and total Stat3 (tStat3) in microglia cell lysates (HMC3) after 8 hours of treatment with the serum-free medium (control condition) and TLQP-21 in the presence or absence of C3a receptor antagonist. (C) Quantification of Stat3 phosphorylation over total Stat3 normalized to tubulin. (n=3 biological replicates; the results are expressed as mean ± SEM). (E) Quantification of IBA1 microglia positive area and (G) intensity per microglia cell in the cerebral cortex of mice after 3 days of treatment with serum-free medium (control condition) and TLQP-21 (4.5 mg/kg). (2 mice/per group, 6-8 cerebral cortex independent areas; the results are expressed as mean ± SEM). Statistical significance was assessed using one-way ANOVA.*P < 0.05, **P < 0.01, ***, P < 0.001, ****P<0,0001.

Importantly, these results were validated *in vivo*. After priming mice with the TLQP-21 for 3 consecutive days, we showed a significant and specific impact on BBB permeability, shown by a significant increase in the accumulation of the fluorescent dye in the brain, when compared with the control condition **(Figure 4F-G)**. Immunohistochemical analyzes of the mice’ brains also showed that TLQP-21 significantly increased IBA1 positive area of microglia cells **(supplementary material, Figure S5B and Figure 5D),** as well as the staining intensity per microglia cell **(supplementary material, Figure S5B and Figure 5E)**.

Collectively, these findings support the functional role of VGF in impacting BBB stability, as well as in the activation of microglia cells during early stages of the brain metastatic cascade, preparing a permissive brain PMN.

#### VGF expression is a poor prognostic marker for breast cancer patients and a predictive factor for brain metastases

Further, we decided to evaluate the clinical impact of VGF on BC prognosis and on BM prediction. For that, we started to study VGF mRNA expression using the Kaplan Meier plotter and VGF protein expression using a large series of primary breast carcinomas. Kaplan-Meier survival analysis revealed that VGF mRNA did not correlate with prognosis (10-year overall survival (OS): hazard ratio (HR)=1.16; p=0.12), when all molecular subtypes of BC were considered. However, VGF mRNA expression was significantly associated with a worse prognosis specifically in triple-negative (10-years OS: HR=1.84, *p*=0.0019) and in HER2 overexpressing (10-years OS: HR=1.93, *p*=0.022) molecular subtypes **(supplementary material, Figure S6)**, which correspond to the molecular subtypes that more frequently metastasize to the brain. Further, we validate this data in a series of primary breast carcinomas, previously characterized by our group (32, 43). Immunohistochemical staining revealed that VGF was preferentially expressed at the cytoplasm of cancer cells and in some cases in the tumor-adjacent stroma, as shown in representative images **(supplementary material, Figure S7)**. Since we were studying the soluble factors secreted by BC cells that could lead to brain PMN remodeling, we considered VGF positivity when there was concomitant expression in cancer cells but also in the tumor-adjacent stroma. We found that BC patients with VGF positive tumors showed significantly lower disease-free survival (DFS) and OS (10-year DFS: *p*=0.009; 10-year OS: *p*=0.011) **(Figure 6A)**. In addition, VGF positivity significantly correlated with poor prognostic markers in BC, such as estrogen receptor (ER) negativity (*p*=0.010), and positive expression for epidermal growth factor receptor (EGFR) (*p*=0.021), metabolic proteins ((CAIX (*p*=0.013) and GLUT1 (*p*=0.004)). More importantly, VGF positive tumors were significantly associated with HER2 overexpression and triple-negative (*p*=0.029) molecular subtypes **(Table 1)**.

**Figure 6.**
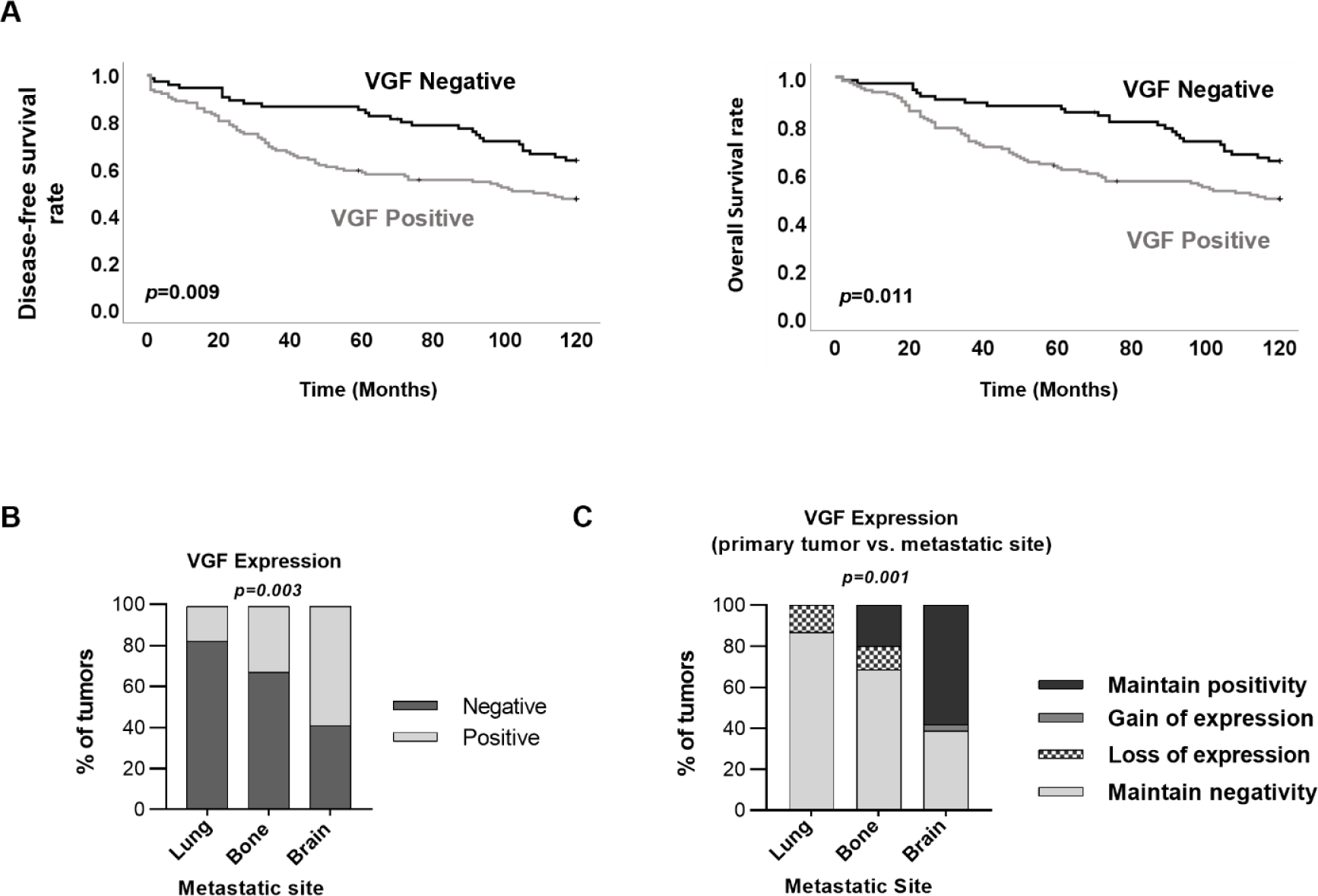
VGF expression associates with a worse prognosis for breast cancer patients and is a predictive factor for brain metastases. (A) Kaplan-Meier plot analysis for overall survival (OS) and disease-free survival (DFS) (OS:*p*=0.011 and DFS:*p*=0.009). (black lines indicate patients with VGF negative tumors; gray lines indicate patients with VGF positive tumors). Survival analyses were compared using the log-rank test. (B) Correlation analysis of VGF expression in primary breast tumors with their corresponding metastatic location. (C) VGF expression in primary breast tumor and its paired metastasis. Chi-square test was used to determine associations between groups and the results were considered statistically significant when the p-value was lower than 0.05.

**Table 1.** Association analysis of VGF expression in primary breast tumors with classical breast cancer poor prognostic factors such as ER (*p*=0.010), EGFR (*p*=0.021), CAIX (*p*=0.013), GLUT1 (p=0.004) and breast cancer molecular subtypes (*p*=0.029). Chi-square test was used to determine associations between groups and the results were considered statistically significant when the p-value was lower than 0.05. ER - Estrogen receptor, PR - Progesterone receptor; EGFR - Epidermal growth factor receptor.

|  |  | VGF Negative |  | VGF Positive |  | p-value |
| --- | --- | --- | --- | --- | --- | --- |
|  |  | frequency | % | frequency | % |  |
| ER | Negative | 15 | 22,4 | 54 | 38,0 | 0,01 |
|  | Positive | 52 | 77,6 | 88 | 62,0 |  |
| PR | Negative | 36 | 52,9 | 88 | 62,0 | ns |
|  | Positive | 32 | 47,1 | 54 | 38,0 |  |
| HER2 | Negative | 64 | 94,1 | 118 | 84,9 | ns |
|  | Positive | 4 | 5,9 | 21 | 15,1 |  |
| EGFR | Negative | 67 | 98,5 | 128 | 89,5 | 0,021 |
|  | Positive | 1 | 1,5 | 15 | 10,5 |  |
| CAIX | Negative | 34 | 53,1 | 49 | 34,8 | 0,013 |
|  | Positive | 30 | 46,9 | 92 | 65,2 |  |
| GLUT1 | Negative | 60 | 92,3 | 105 | 75,5 | 0,004 |
|  | Positive | 5 | 7,7 | 34 | 24,5 |  |
| Molecular subtype | Luminal A-like | 27 | 40,9 | 44 | 32,6 | 0.029 |
|  | Luminal B-like | 26 | 39,4 | 46 | 34,1 |  |
|  | HER 2 overexpressing | 1 | 1,5 | 12 | 8,9 |  |
|  | Triple-Negative | 12 | 18,2 | 33 | 24,4 |  |

Finally, the impact of VGF expression was evaluated on the prediction of the metastatic site. BM series were significantly associated with poor prognostic features, such as high histological grade, age, and molecular subtype (*p*<0.0001) **(supplementary material, Figure S8A-C and Table S2)**. We found that primary breast tumors that metastasized to the brain were associated with poor prognostic factors, such as ER (*p*<0.0001) and progesterone receptor (PR) (*p*=0.002) negative expression, as well as higher proliferative index (assessed by Ki67 labelling) (*p*=0.005), when compared with primary breast tumors that metastasized to other locations (lung and bone) **(supplementary material, Figure S9A)**. As expected, primary breast tumors that metastasized to the brain were mainly from the HER2 overexpression and triple-negative molecular subtypes (*p*<0.0001) **(supplementary material, Figure S9B)**. More importantly, we found that primary tumors with VGF positive expression were significantly associated with an increased and specific propensity to metastasize to the brain (60%), when compared with the metastatic propensity to the lung (20%) or bone (25%) (*p*=0.003) **(Figure 6B)**. Moreover, correlating VGF gain or loss of expression in metastases vs. matched primary breast tumors, we found a significant enrichment of the maintenance of VGF expression in BM **(Figure 6C)** when compared with the other metastatic sites (bone and lung) (*p*=0.001).

## Discussion

BC is the most common cancer among women worldwide, which can metastasize to several parts of the body, including the brain, strongly contributing to cancer-related mortality. A major breakthrough to study the brain metastatic process was the establishment of BC models with a specific tropism to the brain (24).

In this study, we used organotropic MDA-MB-231 cell models to the lung, bone, and brain, which were embedded in collagen type-I, to better mimic the breast microenvironment. Additionally, another brain organotropic cell line transfected with HER2 (231.Brain.HER2) was also included, since its overexpression induces the occurrence of extensive BM in mice (25). Interestingly, we demonstrated that the secretome of both brain organotropic BC cells affect BBB integrity when compared to the secretome of non-brain organotropic cells, which suggest that brain organotropic cells secrete specific factors that play an important role in the remodeling of the brain microenvironment, creating a more permissive site for cancer cell brain colonization. Importantly, the results were always compared to data from non-brain organotropic BC secretomes to control for brain metastatic specificity. Indeed, there are evidence suggesting that factors secreted by cancer cells promote BBB dysfunction by altering the expression and localization of key BBB proteins, namely tight junction proteins. This can lead to increased BBB permeability, allowing cancer cells to enter the brain and potentially form metastases (21, 44–46). In particular, Lu et al. demonstrated that exosomal lncRNA GS1-600G8.5 secreted by brain organotropic BC cells increased *in vitro* BBB permeability, promoting the passage of BC cells across the EC monolayer (45). Moreover, it was recently demonstrated that brain organotropic BC cells secrete CEMIP+ exosomes that can be uptaken by ECs and microglial cells, inducing brain vascular remodeling and inflammation (21).

In addition to a restrictive vascular barrier, the brain is composed of unique resident cells, such as microglia, that play an important role in the immune response after injury. Accordingly, considering cancer progression, microglia cells were described as being responsible for identifying and attacking cancer cells that have invaded the brain (47, 48). Nevertheless, some studies suggest that microglia can also promote the growth and spread of metastatic cells in the brain, contrasting with its tumor surveillance function (35–41). EI Chen et al. demonstrated that NT-3 promote the growth of metastasis by decreasing microglia cell activation (36). In opposition, another study demonstrated that XIST loss in BC cells increased the release of exosomal microRNA-503, which promote *in vitro* anti-inflammatory microglia phenotype (37). Therefore, using the secretome from brain organotropic BC cells, we looked for microglia modifications and found that, in contrast with the specific impact of brain derived secretome on BBB integrity, all secretomes from lung, bone and both brain organotropic cell lines were able to promote microglia activation, as measured by its *in vitro* phagocytic capacity and Stat3 phosphorylation. Considering that *in vitro* microglial cell culture is frequently performed under artificial conditions, which can alter microglial behavior and limit the relevance of experimental findings (49, 50), we also evaluated the impact of the secretomes in microglia using *in vivo* studies. Importantly, we observed that only the secretome of brain organotropic BC cells with HER2 overexpression could promote modifications in microglia of the cerebral cortex, which is the brain region where metastases occur most frequently (3, 51).

Based on these results, we sought to determine the soluble factors responsible for these modifications in the brain PMN. The proteomic analysis revealed that 6 peptides were specifically deregulated in the secretome of both brain organotropic BC models compared to the secretome from lung and bone organotropic cells. VGF emerged as prominent peptide, since it has been suggested to have a role in the brain, both under normal and pathological conditions (43, 52). VGF can be proteolytically processed into multiple bioactive peptides that are involved in brain biological processes related to neuronal growth, differentiation, synaptogenesis, and synaptic plasticity, being extensively studied in several brain disorders (43, 53–57). TLQP-21 is the most common VGF derived peptide, which has been shown to be specifically involved in obesity, diabetes, and neurodegenerative disorders (55, 58–61). In cancer, VGF impact has been underexplored, although it has been regarded as a marker associated with poor prognosis in patients with glioblastoma (62), lung cancer (63), BC featuring a neuroendocrine component (64). However, nothing is known about its role in the context of BM.

Therefore, we went further to evaluate its impact on the brain pre-metastatic remodeling. We demonstrated that TLQP-21 affected BBB integrity and promoted microglia modulation, not only *in vitro* but, most importantly, *in vivo*. Still, the effect of TLQP-21 on microglia modulation was not as pronounced as the secretome of brain organotropic BC cells, suggesting that other factors may also play a role on microglia activation. Interestingly enough, a previous study in Alzheimer’s disease had also shown that TLQP-21 affected the microglia phagocytic capacity (65). Taken together, our findings suggest that VGF is one of the peptides involved in brain remodeling, thus creating a brain pre-metastatic microenvironment, permissive to the success of BMs.

Lastly, the clinical impact of VGF was evaluated in BC samples. We found that VGF expression (both at tumor and stroma) was significantly associated with a poor prognosis for BC patients. Actually, a recent study found that VGF plays a role in conferring resistance to EGFR kinase inhibitors, triggering epithelial-to-mesenchymal transition (EMT) and being associated with a poor prognosis in patients with lung adenocarcinoma (63). It is interesting to note that both lung and BC frequently metastasize to the brain. Furthermore, it has also been shown that VGF have a dual role in glioblastoma, promoting both the survival and self-renewal of glioblastoma stem cells, as well as the proliferation of differentiated glioblastoma cells (62). More importantly, we demonstrated that VGF positive breast tumors specifically correlated with BM, but not to lung or bone metastases, suggesting VGF as a potential predictive biomarker for BM in the BC context.

In summary, we revealed a novel function for VGF in brain remodeling, by promoting BBB disruption and microglia activation. Our data also provide new insights for the role of VGF as a promising prognostic biomarker and therapeutic target for BC patients with an increased risk of developing BM.

## Supporting information

Supplemental Figures andTable

## Acknowledgments

MDA-MB-231 BC organotropic variants were kindly provided by Joan Massagué laboratory. The MDA-MB-231.Brain.HER2 overexpressing cells were kindly provided by Patricia S. Steeǵs laboratory (Center for Cancer Research, National Cancer Institute, Bethesda, MD, USA). We still thank all patients and clinicians for their participation in this study. Dr. Cristiano Souza, Márcia Marques, Vinícius Duval da Silva, Alison Barroso and Daniel Preto compiled patient data from Barretos Cancer Hospital. Jorge Dr. Cameselle-Teijeiro compiled patient data from the Pathology Department, Hospital Xeral-Cíes, Vigo, Spain. The authors acknowledge the i3S Proteomics Scientific Platform. The mass spectrometry technique was performed by Hugo Osório, and the work had the support from the Portuguese Mass Spectrometry Network, integrated in the National Roadmap of Research Infrastructures of Strategic Relevance (ROTEIRO/0028/2013; LISBOA-01-0145-FEDER-022125). The authors acknowledge the assistance of Nuno Mendes from the HEMS core facility at i3S. The i3S Scientific Platform HEMS, is a member of the national infrastructure PPBI—Portuguese Platform of Bioimaging (PPBI-POCI-01-0145-FEDER-022122).

## Statement of author contributions

ASR and RC conceived the structure of the work. RC was involved in all experimental work. RC and IC prepared the secretomes. LS and RL performed the brain EC culture experiments and APS and CG performed the respective data analyses. RC, LS, HF and ASR performed the *in vivo* experiments. CC-M, JL, and CJ compiled patient data from IPOP. FS performed pathology analysis. RC and ASR performed all experimental data analyses. RC and ASR wrote the manuscript. ASR and JP supervised all the work and final-edited the manuscript. All authors have read and agreed to the published version of the manuscript.

## Data Availability

The mass spectrometry proteomics data have been deposited to the ProteomeXchange Consortium via the PRIDE (66) partner repository with the dataset identifier PXD044100. All study data are included in the article and/or **Supplementary Materials and Methods Appendix**.

### Abbreviations

BBB: blood-brain barrier
BC: breast cancer
BCBM: breast cancer brain metastasis
BM: brain metastasis
BSA: bovine serum albumin
C3aRA: complement component 3a-receptor antagonist
C3aR1: complement component 3a receptor-1
CSC: cancer stem cell
DFS: disease free survival
DMEM: Dulbecco’s minimal essential media
EC: Endothelial cell
ECM: extracellular matrix
EGFR: epidermal growth factor receptor
EMT: epithelial-to-mesenchymal transition
ER: estrogen receptor
EVs: extracellular vesicles
FBS: fetal bovine serum
hCM3C/D3: human cerebral microvascular endothelial cell line
HER2: human epidermal growth factor receptor 2
HMC3: human microglial clone 3 cell line
HR: hazard ratio
IFN-y: interferon-y
IP: intraperitoneal injection
IV: intravenous injection
KM: kaplan meier
NaHCO3: sodium bicarbonate
NaOH: sodium hydroxide
NH4Cl: ammonium chloride
OS: overall survival
PBS: phosphate buffered saline
PCA: principal component analysis
PFA: paraformaldehyde
PMN: pre-metastatic niche
PR: progesterone receptor
ROI: region of interest
RT: room temperature
SEM: standard error of the mean
TEER: transendothelial electrical resistance

