## Supplemental Figures andTable for "Deciphering the impact of cancer cell’s secretome and its derived-peptide VGF on breast cancer brain metastasis"

***Ana Sofia Ribeiro**

**Adress: R. Alfredo Allen 208, 4200-135 Porto**

**Phone number: 22 607 4900**

### Conflict of interest statement

The authors have no conflicting interests to disclose.

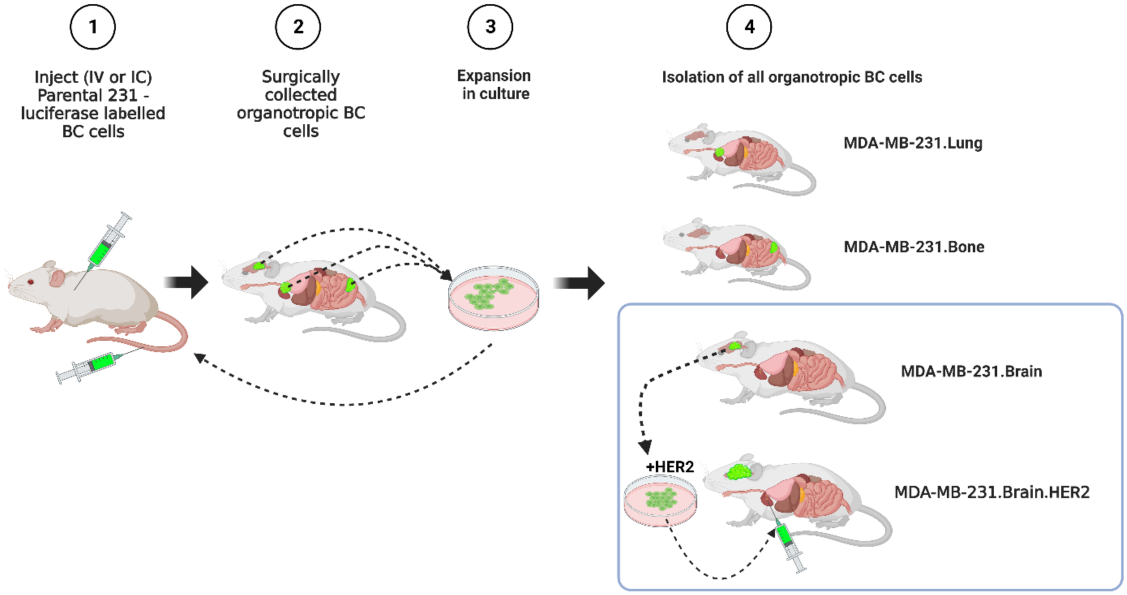

**Figure S1.** Organotropic MDA-MB-231 breast cancer cell model. This model was developed by the injection of the parental 231 cell line into mice by the intracardiac (IC) route and the brain and bone metastases formed were extracted and expanded in culture. These cells were intracardiac re-injected into mice and the process was repeated at least 6 times, originating 231.Brain and 231.Bone organotropic breast cancer cell lines. The same methodology was applied to generate the 231.Lung organotropic breast cell model; however, intravenous (IV) injection in the tail vein was performed. Additionally, the 231.Brain organotropic BC cells were transfected to overexpress HER2 (231.Brain.HER2). Created in Biorender**.**

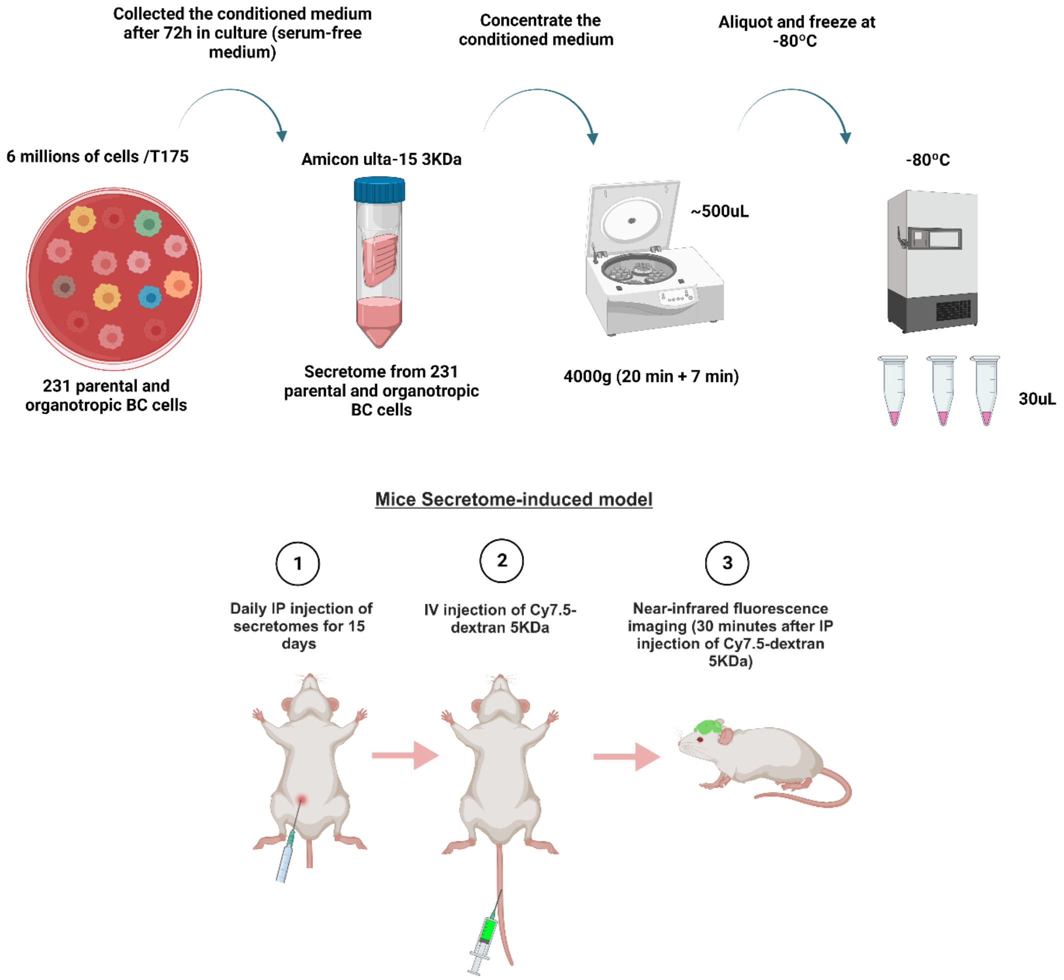

**Figure S2.** Mice secretome-induced model. To understand the impact of the secretomes of parental 231 and their organotropic breast cancer cells *in vivo*, we plated $6x10^6$ cells embedded in collagen type I in a T175 flask. After 24 hours, we added 10 mL of serum-free DMEM for 72 hours. The secretome from the collagen-embedded breast cancer cells was recovered, centrifuged, and concentrated with Amicon® Ultra-15 3KDa Centrifugal Filter (final volume of 500µL). In order to guarantee the integrity of the factors present in the secretomes, we make aliquots of 30 µL and freeze them at -80ºC. The mice with 6–8 weeks of age were pre-treated daily over 15 days with 30 µL of secretome or serum-free DMEM (control condition). At the end of the experiment, near-infrared fluorescence imaging was performed 30 minutes after intravenous injection (IV) of Cy7.5-dextran 5 kDa dye in order to evaluate blood-brain barrier (BBB) integrity. Created in Biorender.

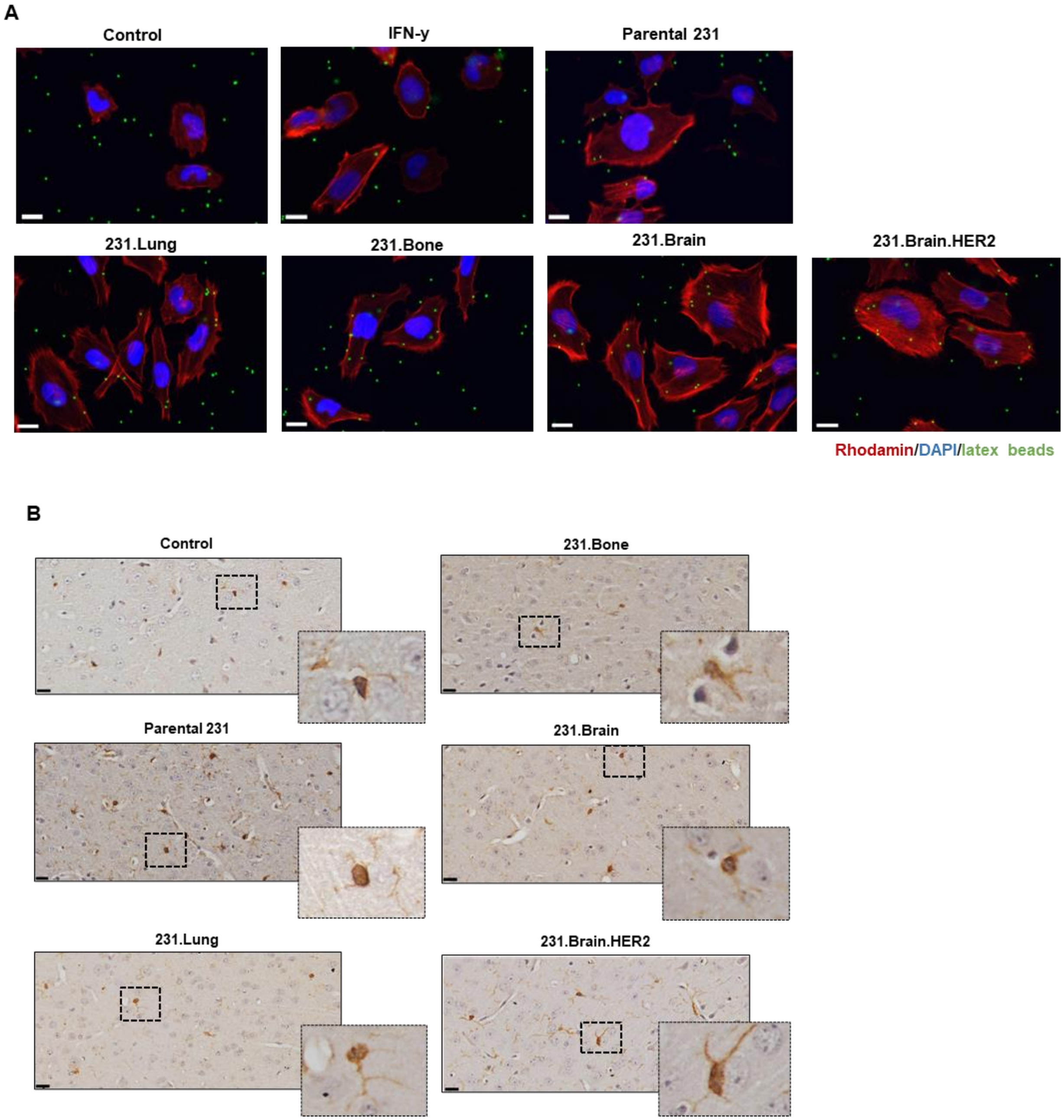

**Figure S3.** The secretome of brain organotropic breast cancer cells impacts microglia activation. (A) Representative confocal images of microglia phagocytic capacity of 1 µm fluorescent latex beads after treatment with serum-free DMEM (control condition), IFN-y (positive control), as well as with the secretomes of parental 231 and their organotropic breast cancer variants for 8 hours. (Scale bar corresponds to 20 µm). (B) Representative images of IHC staining for IBA1 (a marker of microglia cells) in the cerebral cortex of mice after 15 days of treatment with serum-free DMEM (control condition), as well as with the secretomes from the parental 231, and their organotropic breast cancer variants. (Scale bar corresponds to 20 μm).

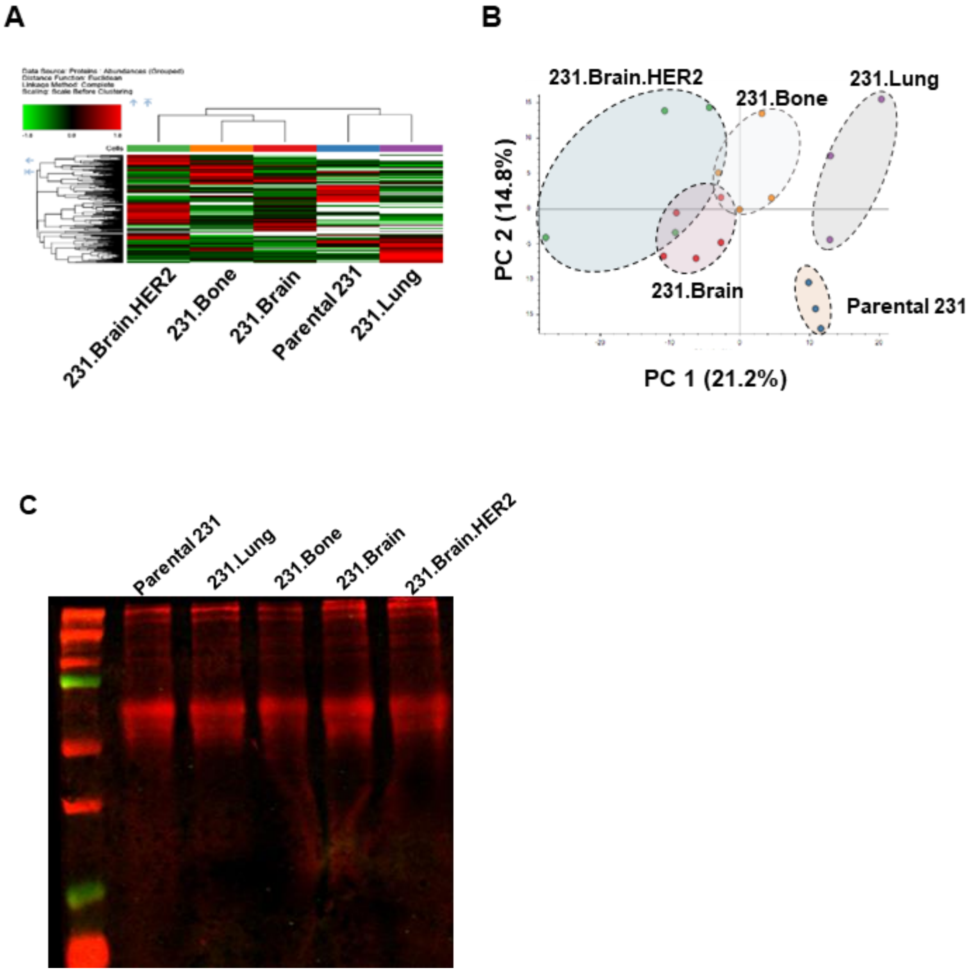

**Figure S4.** Brain organotropic breast cancer cells show a specific secretome signature. (A) Unsupervised hierarchical clustering and (B) Principal component analysis (PCA) of the secretome of parental 231 and their organotropic breast cancer variants. (C) Total protein stain in secretomes detected in the Odyssey® imaging system.

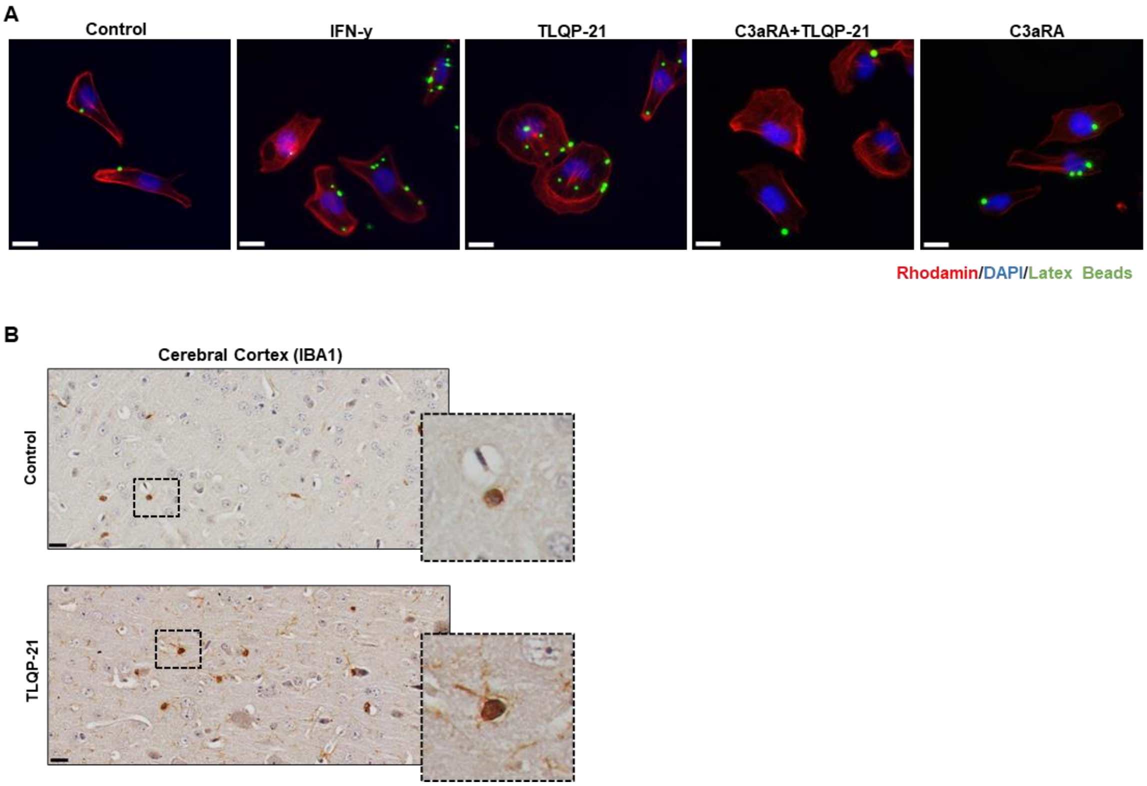

**Figure S5.** VGF impacts microglia activation. (A) Representative confocal images of microglia phagocytic capacity of 1 μm fluorescent latex beads after treatment with serum-free medium (control condition) and TLQP-21 (100nM) in the absence and presence of the C3a receptor antagonist (10μM) for 8 hours. (Scale bar corresponds to 20 μm). (B) Representative IHQ staining for IBA1 in the cerebral cortex of mice after 3 days of treatment with serum-free medium (control condition) and exogenous mouse TLQP-21 (4.5 mg/kg). (Scale bar corresponds to 20 μm).

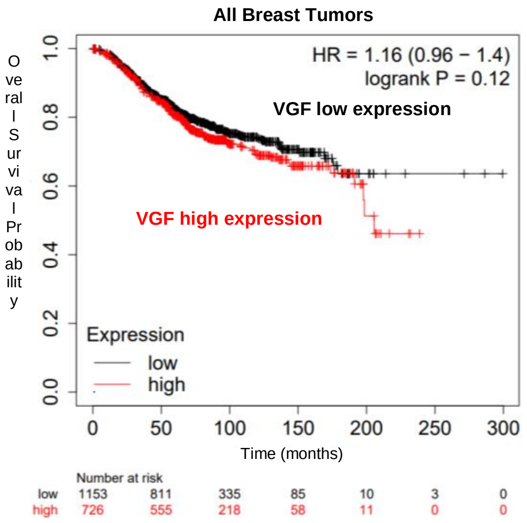
All Breast Tumors
HR = 1.16 (0.96 − 1.4)
logrank P = 0.12
VGF low expression
VGF high expression
Overall Survival Probability
Expression
low
high
Time (months)
Number at risk
low 1153 811 335 85 10 3 0
high 726 555 218 58 11 0 0

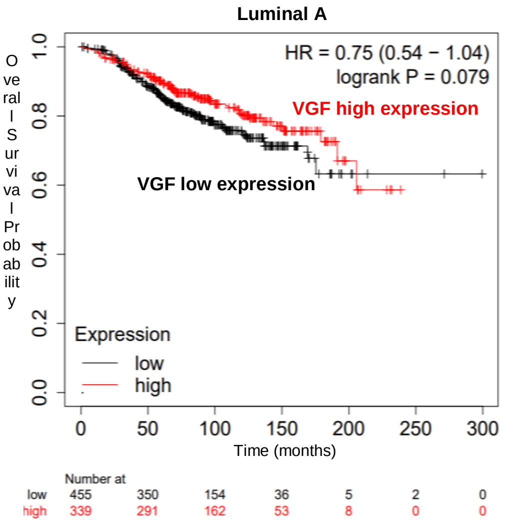
Luminal A
HR = 0.75 (0.54 − 1.04)
logrank P = 0.079
VGF high expression
VGF low expression
Overall Survival Probability
Expression
low
high
Time (months)
Number at
low 455 350 154 36 5 2 0
high 339 291 162 53 8 0 0

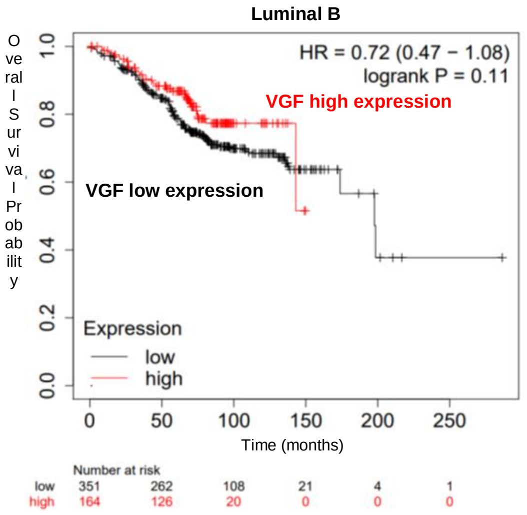
Luminal B
HR = 0.72 (0.47 − 1.08)
logrank P = 0.11
VGF high expression
VGF low expression
Overall Survival Probability
Expression
low
high
Time (months)
Number at risk
low 351 262 108 21 4 1
high 164 126 20 0 0 0

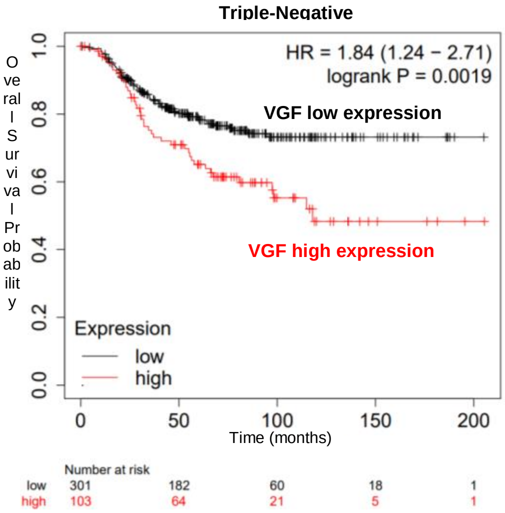
Triple-Negative
HR = 1.84 (1.24 − 2.71)
logrank P = 0.0019
VGF low expression
VGF high expression
Overall Survival Probability
Expression
low
high
Time (months)
Number at risk
low 301 182 60 18 1
high 103 64 21 5 1

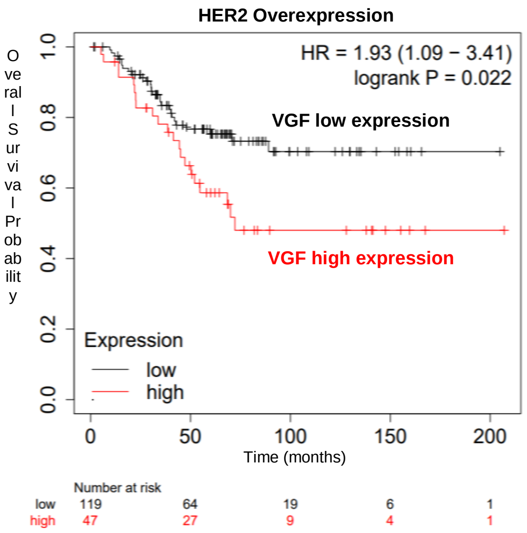
HER2 Overexpression
HR = 1.93 (1.09 − 3.41)
logrank P = 0.022
VGF low expression
VGF high expression
Overall Survival Probability
Expression
low
high
Time (months)
Number at risk
low 119 64 19 6 1
high 47 27 9 4 1

**Figure S6.** Kaplan-Meier survival curves of breast cancer patients based on VGF mRNA expression. (black lines indicate patients with VGF low expression; red lines indicate patients with VGF high expression). Kaplan-Meier plots comprising all breast tumors (Overall-Survival (OS): hazard ratio (HR) =1.16, p=0.12)), Luminal A (OS: HR=0.75, p=0.079), Luminal B (OS: HR =0.72, p=0.11), triple-negative (OS: HR=1.84, p=0.0019), and HER2 overexpressing breast tumors (OS: HR=1.93, p=0.022).

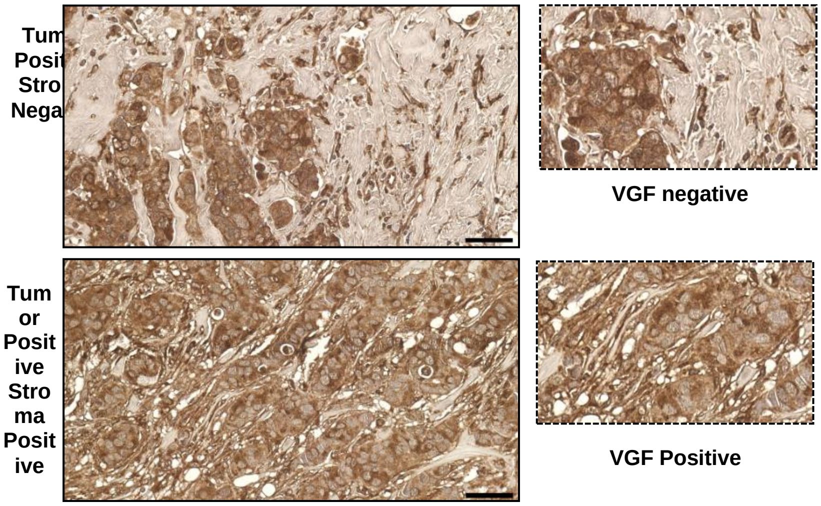

**Figure S7.** VGF expression associates with a worse prognosis for breast cancer patients and is a predictive factor for brain metastases. (A) Representative images of VGF expression in tumor (VGF negative) and co-expression in tumor and stroma (VGF positive) in primary breast tumors. (Scale bar corresponds to 50 μm).

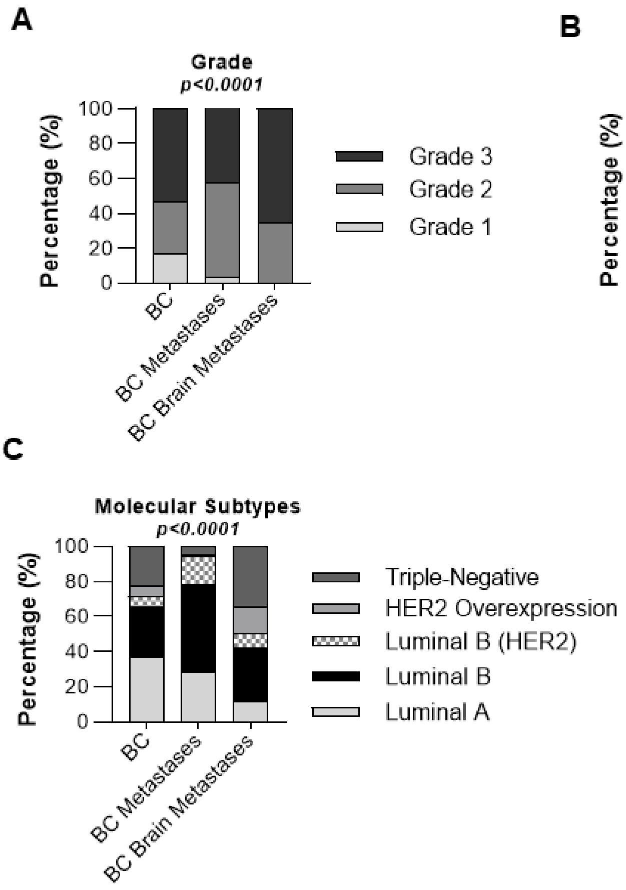

**Figure S8.** Association of the different breast cancer cohorts with clinico-pathological features. Correlation analysis between different BC cohorts with (A) histological grade (p<0.0001), (B) age (p<0.001), and (C) molecular subtypes (p<0.0001). The chi-square test was used to determine associations between groups and the results were considered statistically significant when the p-value was lower than 0.05. (BC – primary breast tumors, BC metastases – primary breast tumors with metastases, and BC Brain Metastasis – primary breast tumors with brain metastases only).

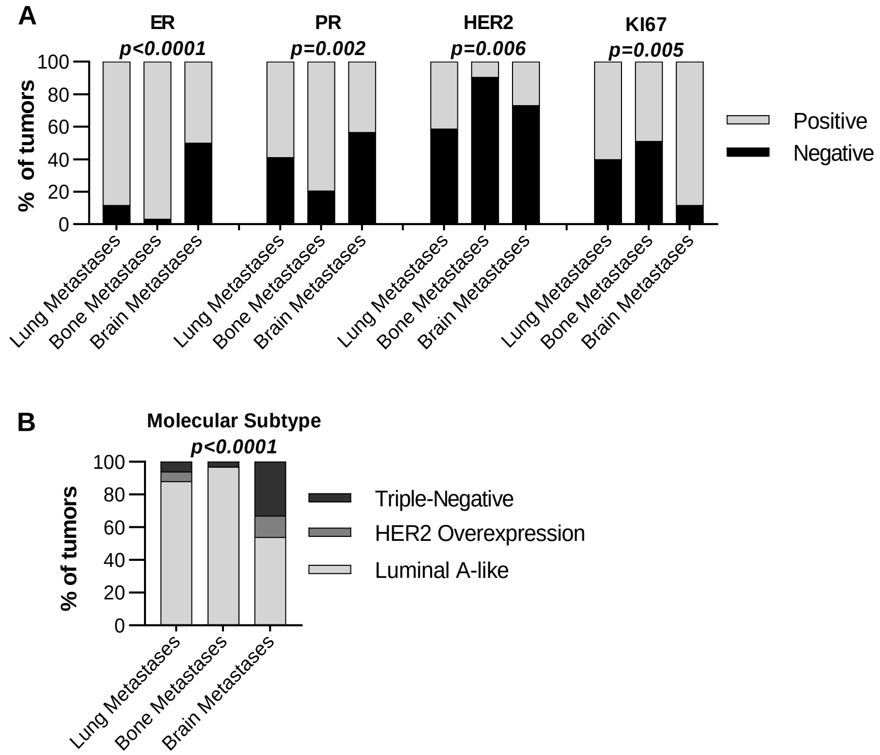

**Fig S9.** Association between primary breast tumors that metastasize to the lung, bone, and brain with classical poor prognostic factors and molecular subtypes of breast cancer. (A) Correlation analysis of primary breast tumors that metastasize to the lung, bone, and brain with classical breast cancer poor prognostic factors such as ER (p<0.0001), PR (p=0.002), and Ki67 (p=0.005). (B) Correlation analysis of primary breast tumors that metastasize to the lung, bone, and brain with breast cancer molecular subtypes (p<0.0001). The chi-square test was used to determine associations between groups and the results were considered statistically significant when the p-value was lower than 0.05.

**Table S1.** Characterization of the different breast cancer cohorts.

| DATABASE | BC | BC METASTASIS | BRAIN METASTASIS |
|---|---|---|---|
| COUNTRY | SPAIN | PORTUGAL | BRASIL |
| HOSPITAL | CHUVI | IPO-P | Barretos |
| ETHICAL PROTOCOL NUMBER/APPROVAL | | | 2.777.372 |
| TYPE BREAST TUMORS | Primary | Primary and Metastasis | Primary and Metastasis |
| METASTASIS | No information | Bone, Lung, and Brain | Brain |
| NUMBER | 218 | 77 | 27 |
| TYPE | retrospective | retrospective | retrospective |
| DATE | 1978-1992 | 1996-2016 | 2012-2018 |
| Age at diagnosis | 59,98 ± 12,637 (32-90) | 51,07 ± 12,35 (28-76) | 49,04 ± 12,73 (27-71) |
| Follow-up duration | 120.0 | 120.0 | 120.0 |
| Disease-free survival (DFS; months) | 84,58 ± 44,7 (1-120) | 73,18 ± 52,37 (0-120) | 28,2 ± 28,23 (0-110) |
| Overall Survival (OS; months) | 88,25 ± 341,4 (2-120) | 87,34 ± 34,58 (18-120) | 53,29 ± 33,98 (6-120) |

**Table S2.** Association of the different breast cohorts with clinico-pathological features.*

| | | BC | | BC METASTASIS | | BRAIN METASTASIS | | |
|---|---|---|---|---|---|---|---|---|
| | | frequency | % | frequency | % | frequency | % | p-value |
| QT_Neo | No | - | - | 74 | 90,2 | 16 | 59.3 | - |
| | Yes | - | - | 8 | 9,8 | 11 | 40.7 | |
| Grade | 1 | 37 | 17,5 | 3 | 3,7 | 0 | 0 | <0,0001 |
| | 2 | 62 | 29,2 | 44 | 53,7 | 9 | 34,6 | |
| | 3 | 113 | 53,3 | 35 | 42,7 | 17 | 65,4 | |
| | missing | 6 | | 8 | | 1 | | |
| Age | < 50 years old | 84 | 24,2 | 42 | 50,6 | 16 | 59,3 | <0,0001 |
| | > 50 years old | 263 | 75,8 | 41 | 49,4 | 11 | 40,7 | |
| Lymph Node | Negative | 63 | 36,8 | 19 | 23,5 | 8 | 29.6 | ns |
| | Positive | 108 | 63,2 | 62 | 76,5 | 18 | 66.7 | |
| | missing | 47 | | 7 | | 1 | 3.7 | |
| Metastasis at Diagnosis | Negative | - | - | 71 | 85,5 | 21 | 77.8 | ns |
| | Positive | - | - | 12 | 14,5 | 6 | 22.2 | |
| | Undefined | - | - | 0 | | | 0.0 | |
| Molecular subtype | Luminal A-like | 76 | 36,7 | 24 | 28,9 | 3 | 11,5 | <0,0001 |
| | Luminal B-like (HER2 negative) | 59 | 28,5 | 41 | 49,4 | 8 | 30,8 | |
| | Luminal B-like (HER2 positive) | 13 | 6,3 | 13 | 15,7 | 2 | 7,7 | |
| | HER 2 overexpressing | 13 | 6,3 | 1 | 1,2 | 4 | 15,4 | |
| | Triple-Negative | 46 | 22,2 | 4 | 4,8 | 9 | 34,6 | |
| | Undefined | 11 | | 7 | | 1 | | |

*Association of the different breast cancer cohorts (BC, BC metastasis, and Brain metastasis) with QT_Neo, grade, age, lymph node, metastasis at diagnosis, and molecular subtypes. The chi-square test was used to determine associations between groups and the results were considered statistically significant when the p-value was lower than 0.05. (BC – primary breast tumors, BC metastases – primary breast tumors with metastases, and BC Brain Metastasis – primary breast tumors with brain metastases only).
