## Supplemental Materials and Methods for "Deciphering the impact of cancer cell’s secretome and its derived-peptide VGF on breast cancer brain metastasis"

***Ana Sofia Ribeiro**

**Adress: R. Alfredo Allen 208, 4200-135 Porto**

**Phone number: 22 607 4900**

### Conflict of interest statement

The authors have no conflicting interests to disclose.

### Supplementary Materials and Methods

#### Cell Culture

The parental BC cell line MDA-MB-231 (parental 231) was obtained from ATCC (American Type Culture Collection, Manassas, VA, USA), while its organotropic BC variants lung (231.lung) (1), bone (231.Bone) (2), brain (231.Brain) (3) and brain with HER2 overexpression (231.Brain.HER2) (4) clones were obtained from J Massagué (MSKCC) and P. Steeg (NCI) (**Supplementary Figure S1**). The five cell lines were maintained in Dulbecco's minimal essential media (DMEM), supplemented with 10% fetal bovine serum (FBS), and with 1% antibiotic solution penicillin/streptomycin (all reagents from Invitrogen, Carlsbad, CA, USA). The human cerebral microvascular endothelial cell line (hCMEC/D3), used as an *in vitro* model of the BBB (5), was kindly provided by Pierre-Olivier Couraud (Institute Cochin, Université René Descartes, Paris, France). This cell line was cultured in Endothelial Basal Medium-2 (EBM-2; Lonza, Walkersville, MD) supplemented with 1% chemically defined lipid concentrate (Invitrogen, Inchinnan Business Park, UK), 1 ng/ml basic fibroblast growth factor (bFGF; Sigma-Aldrich), 10 mM 4-(2-hydroxyethyl)-1-piperazineethanesulfonic acid (Lonza), 1.4 µM hydrocortisone (Sigma-Aldrich), 5 µg/ml ascorbic acid (Sigma-Aldrich), 1% penicillin-streptomycin (Gibco), and 5% FBS (Invitrogen) on culture plates coated with collagen Type I (R&D Systems, Inc., Minneapolis, MN). The medium was changed every 2 days until the cells reached confluence. The human microglial clone 3 cell line (HMC3) was purchased from ATCC (ATCC® CRL-3304™) and grown in Eagle's Minimum Essential Medium (EMEM; ATCC, VA, USA), supplemented with 10% FBS and

1% penicillin/streptomycin (all reagents from Invitrogen, Carlsbad, CA, USA). All the cell lines were cultured at 37ºC in a humidified atmosphere with 5% CO2. Cells were used in experiments upon reaching 70–80% confluence.

**Secretomes Preparation**

Using the parental 231 and their organotropic BC variants to the lung, bone, and brain (without or with HER2 overexpression), we plated (3x$10^5$ cells) in a 6 well-plate, which were subsequently embedded in collagen type I (Merck) to mimic the breast ECM. After 24 hours, we added 1 mL serum-free DMEM to the 3D cultures and left them for 72 hours. The secretomes were recovered, centrifuged (to eliminate cell debris), and used to perform all *in vitro* experiments.

In order to study the impact of the secretomes *in vivo*, we plated BC cells (6x$10^6$ cells) embedded in collagen type I in a T175 flask. After 24 hours, we added 10 mL serum-free DMEM and left them for 72 hours. The secretomes from the collagen-embedded BC cells were recovered, centrifuged, and concentrated with Amicon® Ultra-15 3KDa Centrifugal Filter (Millipore) until a final volume of 500µL. In order to guarantee the integrity of the factors present in the secretomes, we made aliquots of 30µL and freeze them at -80ºC (**Supplementary Figure S3**).

The collagen type I from the rat tail (final concentration of 0.2 mg/mL) was mixed with phosphate buffered saline (PBS), serum-free DMEM, sodium bicarbonate (NaHCO3), and sodium hydroxide (NaOH), in ice-cold conditions.

***In vitro* evaluation of brain endothelial cell monolayer integrity**

Cell monolayer integrity was determined by measuring the transendothelial flux of fluorescent 4kDa macromolecule across the endothelial cells (ECs) and the transendothelial electrical resistance (TEER) as previously described (6). Briefly, hCMEC/D3 were grown on collagen type I (R&D Systems, Inc., Minneapolis, USA) coated 12 mm Transwell filters (Costar, Corning, NY, USA) at a density of $5.4x10^4$ cell/$cm^2$, and were maintained for 7 days in a 5% CO2 incubator at 37°C in regular culture medium for endothelial monolayer formation. On the seventh day, the culture medium from the upper compartment was removed and replaced by the secretomes from parental 231and their organotropic BC cells for 24 hours. ECs exposed to fresh serum-free DMEM were used as control.

In addition, exogenous human purified TLQP-21 peptide at a concentration of 100 nM (Bachem, Bubendorf, Switzerland) was added to the serum-free DMEM. To determine whether TLQP-21 was involved in EC barrier dysfunction, the complement component 3a-receptor antagonist (C3aRA; Sigma-Aldrich; 204881) was added to ECs at concentration of 10µM during incubation with the secretome of 231.Brain.HER2 overexpression or exogenous TLQP-21, respectively. In order to evaluate the permeability of EC monolayer, 1mg/mL fluorescein isothiocyanate (4kDa FITC; Sigma-Aldrich) was added to the apical side of the transwell containing ECs monolayers, diluted in serum-free DMEM or in secretomes of parental 231 and their organotropic BC variants. Samples (50µL) were collected from the basal chamber (and replaced by the corresponding medium) at 20 minutes intervals for 180 minutes after treatments. The fluorescence of samples was measured in a microplate reader (Biotek, Synergy HT, Winooski, VT) and plotted against time. Permeability was determined from linear slope changes after treatments. TEER of ECs monolayers was measured using an STX-2 electrode coupled to an EVOM resistance meter (World Precision Instruments, Hertfordshire, UK). TEER readings of cell-free inserts were subtracted from the values obtained with cells, and results were expressed as a percentage of control. TEER measurements were performed 24 hours after the replacement of the cell culture medium by the above-mentioned secretomes or serum-free DMEM (in the control condition). Only ECs monolayers with a TEER above 50 Ωcm2 were used in subsequent experiments and were obtained by subtracting the TEER of empty inserts giving the specific TEER of ECs monolayers.

***In vitro* microglial phagocytosis assay**

HMC3 cells were seeded (2.5x$10^4$ cells) in a 12-well for 24 hours. Then, we treated microglia cells with the secretomes from parental 231and their organotropic BC cells for 8 hours. Microglia cells exposed to serum-free DMEM were used as control. In addition, we treated microglia cells with 0.2 mg/mL Interferon-γ (IFN-y; Citomed; 285-IF-100) as a positive control.

Exogenous TLQP-21 peptide at a concentration of 100 nM (Bachem, Bubendorf, Switzerland) was added to serum-free DMEM. To determine whether TLQP-21 was involved in microglia phagocytic capacity, the C3aRA was added to microglia cells at concentration of 10 μM during incubation with exogenous TLQP-21 for 8 hours. To evaluate phagocytic capacity, microglia cells were incubated with 0.0025% (w/w) 1 μm fluorescent latex beads for 75 minutes at 37 °C. At the end of the incubation period, cells were fixed with freshly prepared 4 % (w/v) paraformaldehyde (PFA) in PBS for 10 minutes, and then washed with PBS four times to remove unbound beads. Total microglia cells and the number of ingested beads per cell were counted per condition using ImageJ software in order to determine the percentage of phagocytic cells (7). We only consider microglia phagocytic capacity when we found more than 5 fluorescent latex beads per cell.

**Mice secretome induced-model**

In this study, we optimized an *in vivo* secretome induced-model to study the impact of the secretome of parental 231 and of their organotropic BC cells on the remodelling of BBB integrity and microglia modulation. Additionally, we also evaluated the impact of mouse TLQP-21 *in vivo* (8, 9).

In detail, 2 to 3 Female N:NIH(S)II-nu/nu mice per condition, 6–8 weeks of age were pre-treated via intraperitoneal injection (IP) with 30 μL serum-free DMEM (control condition), 30 μL secretome from 231 parental and their organotropic BC variants as well as 30 μL of MCF7 secretome, as a negative control (non-metastatic BC cells) for 15 consecutive days.

In order to understand the impact of TLQP-21 on BBB integrity we pre-treated the animal (4 Female N:NIH(S)II-nu/nu mice per condition, 6–8 weeks) with mouse TLQP-21 (4.5 mg/kg; Sigma-Aldrich T1581) for 3 days. Mice follow-up was performed as previously described (10).

Near-infrared fluorescence imaging was performed 30 minutes after intravenous injection (IV) of Cy7.5-dextran 5 kDa (1.5 mg/kg; Nancocs) dye to evaluate BBB permeability (**Supplementary Figure S2**). The analysis of bioluminescent images was conducted using the Living Image software. To quantify the bioluminescent signal, a region of interest (ROI) was delineated around the lesions.

At the end of the experiment, mice were anesthetized with sodium pentobarbital (80 mg/kg, i.p., Sigma-Aldrich) and transcardially perfused with 10 mL of 0.05 mol/L sodium citrate in 1% PFA (pH 4.2, 37°C), followed by 20 mL of 4% PFA in 0.01 mol/L PBS, pH 7.4. Brains were removed and embedded in paraffin for further immunofluorescence and immunohistochemistry analyses.

The mice were housed, bred, and maintained at the i3S animal house in a pathogen-free environment under controlled conditions of light and humidity. All the experiments were conducted with the application of the 3Rs (replacement, reduction, and refinement) (JP_2016_02 Project, animal ethics committee, and animal welfare body of i3S) (11).

**Immunofluorescence assay**

Cells were cultured on glass coverslips, fixed with 4% PFA for 20 minutes, treated with ammonium chloride (NH4Cl) (50 mM) for 10 minutes, washed with PBS, and permeabilized with 0.1% Triton X-100 in PBS for 5 minutes, at room temperature (RT). Non-specific binding was blocked by treatment with 5% bovine serum albumin (BSA) in PBS, for 30 minutes at RT. Cells were then stained with Rhodamine Phalloidin (1:200, Life Technologies), for 30 minutes in order to visualize F-actin, or with anti-β-catenin (1:150), and anti-ZO-1 (1:100) overnight at 4ºC followed by conjugated goat anti-mouse and anti-rabbit secondary IgG (Dako

Cytomation, Carpinteria, CA, USA), for 1 hour at RT. After another wash with PBS, each sample was mounted with Vectashield (Vector Laboratories, Inc., Burlingame, CA, USA) containing 4′,6-diamidino-2-phenylindole (DAPI) and visualized with Zeiss LSM 710 confocal microscope (Carl Zeiss AG, Jena, Germany). Brain tissue was postfixed in 4% PFA for 24 hours at RT and transferred to 30% sucrose in 0.01 mol/L PBS, pH 7.4, for at least 24 hours at 4°C. Next, coronal sections (12 μm) were cut on a cryostat (Leica CM3050S, Nussloch, Germany), mounted directly onto super-frost microscope slides (Thermo Scientific, Menzel GmbH & Co KG, Braunschweig, Germany) and stored at - 80°C until further use. Then, slices were rinsed in 0.01 mol/L PBS, blocked with 5% BSA in 0.01 mol/L PBS for 1 hour at RT, and incubated overnight at 4°C with rabbit anti-collagen IV (1:200; Abcam) and goat anti-albumin (1:2000; Bethyl Laboratories, Inc, Montgomery, TX, USA) antibodies. Afterward, slices were incubated with Alexa Fluor 488 and Alexa Fluor 594 secondary antibodies (1:200; Invitrogen) for 1 hour at RT, followed by nuclei staining with 5 μg/mL Hoechst 33342 (Sigma-Aldrich) for 5 minutes at RT in the dark. Finally, slices were mounted with Dako fluorescence medium (Dako North America), and images were recorded using a LSM 710 Meta Confocal microscope (Carl Zeiss).

**Western Blot**

Cells were lysed with PBS containing 1% Nonidet/P40 (Sigma-Aldrich, Darmstad, Germany), 1% Triton X100 (Sigma-Aldrich, Darmstad, Germany), 1:7 Protease Inhibitors Cocktail (Roche Diagnostics GmbH, Mannheim, Germany), and 1:100

Phosphatase inhibitor (Sigma-Aldrich, Darmstad, Germany).

Protein concentration (cell lysates or secretomes) was determined using the Bio-Rad protein assay (BIO-RAD, Richmond, CA, USA) and samples were loaded into a 10% polyacrylamide gel and transferred onto a nitrocellulose membrane (GE Healthcare Life Sciences, Sheffield, UK) at 100 V for 90 minutes. Membranes were blocked in 0.5% Tween 5% nonfat dry milk for 1 hour, stained with specific primary antibodies overnight at 4º C, and with secondary antibodies for 1 hour at RT. Detection was assessed using the ECL Chemiluminescence detection kit (BIO-RAD, Richmond, CA, USA). Primary antibodies were anti-VGF (dilution 2 µg/ml, Abcam), anti-phospho-Stat3 (Tyr705) (dilution 1:2000, Cell Signaling), anti-Stat3 (dilution 1:1000, Cell Signaling) and anti-tubulin (dilution 1:1000, Sigma-Aldrich, Darmstad, Germany). Peroxidase-conjugated secondary antibodies (anti-rabbit and anti-mouse) were purchased from Santa Cruz Biotechnology Inc. (Heidelberg, Germany).

To visualize proteins present in secretomes following protein electrophoresis and transfer, we employed the Revert™ 700 Total Protein Stain method, a well-established protocol designed for protein detection and visualization.

First, the membrane is rinsed with ultrapure water to prepare it for staining. Subsequently, the membrane is treated with the Revert 700 Total Protein Stain, allowing the proteins to be distinctly marked for visualization. Following staining, a thorough rinse with a wash solution is performed to remove any excess stain. A final rinse with ultrapure water ensures a clean background for accurate imaging.

To capture images of the membrane, we used an Odyssey® Imaging System with the 700 and 800 nm channels. Adjust the scan intensity or acquisition time to prevent saturation.

**Proteomic analysis**

Proteins were reduced and alkylated with 100 mM Tris pH 8.5, 1% sodium deoxycholate, 10 mM tris (2-carboxyethyl) phosphine (TCEP), and 40 mM chloroacetamide for 10 minutes at 95ºC at 1000 rpm (Thermomixer, Eppendorf). Next, 100 µg of protein were processed for proteomics analysis following the solid-phase-enhanced sample-preparation (SP3) protocol as described by Hughes et al (12). Enzymatic digestion was performed with Trypsin/LysC (2 µg) overnight at 37ºC at 1000 rpm. The resulting peptides were cleaned-up and desalted with C18 micro columns and further quantified. Protein identification was performed by nanoLC-MS/MS. This equipment is composed of an Ultimate 3000 liquid chromatography system coupled to a Q-Exactive Hybrid Quadrupole-Orbitrap mass spectrometer (Thermo Scientific, Bremen, Germany). Peptides were loaded onto a trapping cartridge (Acclaim PepMap C18 100Å, 5 mm x 300 µm i.d., 160454, Thermo Scientific) in a mobile phase of 2% ACN, 0.1% FA at 10 µL/min. After 3 minutes loading, the trap column was switched in-line to a 50 cm by 75µm inner diameter EASY-Spray column (ES803, PepMap RSLC, C18, 2 µm, Thermo Scientific, Bremen, Germany) at 300 nL/minute. Separation was generated by mixing A: 0.1% FA, and B: 80% ACN, with the following gradient: 5 minutes (2.5% B to 10% B), 120 minutes (10% B to 30% B), 20 minutes (30% B to 50% B), 5 minutes (50% B to 99% B) and 10 minutes (hold 99% B). Subsequently, the column was equilibrated with 2.5% B for 17 minutes. Data acquisition was controlled by Xcalibur 4.0 and Tune 2.9 software (Thermo Scientific, Bremen, Germany). The mass spectrometer was operated in data-dependent (dd) positive acquisition mode alternating between a full scan (m/z 380-1580) and subsequent HCD MS/MS of the 10 most intense peaks from the full scan (normalized collision energy of 27%). ESI spray voltage was 1.9 kV. Global settings: use lock masses best (m/z445.12003), lock mass injection Full MS, chrom. peak width (FWHM) 15s. Full scan settings: 70k resolution (m/z 200), AGC target $3e^{6}$, maximum injection time 120 ms. dd settings: minimum AGC target 8e3, intensity threshold 7.3e4, charge exclusion: unassigned, 1, 8, >8, peptide match preferred, exclude isotopes on, dynamic exclusion 45s. MS2 settings: microscans 1, resolution 35k (m/z 200), AGC target $2e^{5}$, maximum injection time 110 ms, isolation window 2.0 m/z, isolation offset 0.0 m/z, spectrum data type profile. The raw data was processed using the Proteome Discoverer software (Thermo Scientific) and searched against the UniProt database for Homo sapiens Proteome. The Sequest HT search engine was used to identify tryptic peptides. The ion mass tolerance was 10 ppm for precursor ions and 0.02 Da for-fragment ions. The maximum allowed for missing cleavage sites was set to 2. Cysteine carbamidomethylation was defined as constant modification. Methionine oxidation and protein N-terminus acetylation were defined as variable modifications. Peptide confidence was set to high. The processing node Percolator was enabled with the following settings: maximum delta Cn 0.05; decoy database search target FDR 1%, validation based on qvalue. Imputation of missing values was performed only when a peptide was detected in at least half (two) of the replicates analyzed. Quantitative proteomic evaluation was performed by pairwise comparisons of all detected peptides and the median ratio was used for protein level comparison. Significance assessment was performed using the background-based ANOVA method implemented in Proteome Discoverer 2.4.1.15 and multiple comparison adjustment of the p values was performed. A minimum of two unique peptides and quantitation in at least half of the samples were required for further evaluation of each protein. The Venn diagrams were generated with the web tool available at bioinformatics http://www.ehbio.com/test/venn/.

**Kaplan Meier plotter survival analysis for VGF mRNA**

Kaplan Meier (KM) plotter online survival analysis tool (https://kmplot.com) was used to assess the impact of VGF mRNA levels on overall survival (OS) in BC patients, being the probe sets the following: 205586_x_at. The “Auto select best cut-off” option was used since it computes all possible cut-off values and determines the best-performing threshold for the survival analysis. Using these parameters, the analysis was carried out on 1879 patients for OS.

**Primary Breast Cancer Series**

A series of 218 primary breast carcinomas diagnosed between 1978–1992, were retrieved from the Pathology Department, Hospital Xeral-Cíes, Vigo, Spain. Patient follow-up information was available for the 218 (**Supplementary Table S1 and S2**), with a maximum follow-up of 120 months after diagnosis. The tumors were characterized for clinical and pathological features, as previously described (13). The disease-free survival (DFS) interval was defined as the time from diagnosis to the date of breast cancer-derived relapse (DFS: mean with 95% confidence interval (CI) of 85.5 +/− 2.1 months), whereas overall survival (OS) was considered as the number of months from diagnosis to the disease-related death (OS: mean with 95% CI of 90.0 +/− 2.0 months). BC patients followed the adequate protocols for chemotherapy, radiotherapy, and hormone therapy given at that time. All patients were treated with adjuvant chemotherapy, which consisted of a protocol of six cycles of cyclophosphamide, methotrexate, and fluorouracil. Patients with ER-positive tumors were treated with hormonal therapy, which was carried out exclusively with tamoxifen. Patients with HER2-overexpressing carcinomas were not treated with specific targeted therapy (trastuzumab). No neo-adjuvant treatment was used in the patients included in this series. The present study was conducted under the national regulative law for the usage of biological specimens from tumor banks, where the samples are exclusively available for research purposes in the case of retrospective studies. All analyses were performed according to the reporting recommendations for tumor MARKer prognostic studies (REMARK) recommendations for prognostic and tumor marker studies.

**Breast Cancer Metastases series**

We used two complementary paired primary and metastatic BC series

(**Supplementary Table S1 and S2**). The first one was retrospectively collected from the archives of the Department of Pathology of the Portuguese Oncology Institute of Porto (IPO Porto) and includes primary breast tumors that metastasized to the lung (n=17), bone (n=56), and brain (n=4). The tumors corresponded to biopsy and/or surgical resections, and were formalin-fixed and parafin-embedded as per routine diagnostics, being evaluated by a dedicated Breast Pathologist. All patients were treated by the same multidisciplinary team of professionals in the Breast Cancer Clinic of IPO Porto. The second one is an exclusive BCBM series with paired primary breast tumors and their corresponding brain metastasis, which was retrospectively collected from Barretos Cancer Hospital, Brazil. Breast carcinomas were classified considering European Society for Medical Oncology guidelines into luminal A (estrogen receptor (ER)/progesteron receptor (PR) positive, HER2 negative, and Ki67 low), luminal B (ER/PR positive, HER2 and/or Ki67 high), HER2-overexpressing (ER/PR negative, HER2 positive), and triple-negative (ER/PR/HER2 negative) carcinomas. Metastases present in BC patients were collected after surgical resection or postmortem, preserved and fixed in formalin, included in paraffin, and carefully evaluated by neuropathologists. Clinical and pathological features were retrieved for this study from clinical charts. At diagnosis, the majority of patients were stage IV (26.8%, 15/56), followed by stage III (19.6%, 11/56), stage II (26.7%, 15/56), stage I (14.3%, 8/56), and for 7 patients we did not have information on the stage. Information regarding prior metastatic dissemination to the brain was not complete and it was not used in the analysis. Whenever data were available, patients were grouped into intrinsic molecular subtypes: luminal (positive for ER and/or progesterone receptor (PR), HER2-negative), HER2-positive (HER2-positive/amplified, negative or positive for ER and PgR), and triple-negative (ER-negative and PgR-negative, HER-2-negative). Patient follow-up information was available, for those who were diagnosed between 2007–2008 (Barretos Cancer Hospital), with a maximum follow-up of 223 months after diagnosis. These BC patients followed the established protocols of chemotherapy, radiotherapy, and hormone therapy at that time. No neo-adjuvant treatment had been used in the patients included in this series.

The present study was conducted with the approval of the Ethical Commission from both cancer centers, under the national regulative law for the usage of biological specimens from tumor banks, where the samples are exclusively available for research purposes in retrospective studies (Ethical approvals: Portuguese Oncology Institute of Porto (CES. 64/023) and Barretos Cancer Hospital/Fundação Pio XII (2-777-372)).

**Immunohistochemistry**

Immunohistochemistry for IBA1 and VGF were performed in 3 µm sections. Slides were placed in a Clear-Rite bath (Thermo Fisher Scientific, Waltham, MA, USA), rehydrated through a descending series of ethanol washes, and finally placed in distilled water. Epitope exposure was performed for 15 minutes (IBA1 antibody) or 30 minutes (VGF antibody) at 95 ◦C with citrate buffer (ThermoScientific, Freemont, CA, USA). The following antibodies were used: IBA1 (Thermo Fisher

Scientific; GT10312; 1:200) and VGF (Abcam;115609; 5μg/mL). Primary antibodies were detected using the horseradish peroxidase polymer (Cytomation Envision System HRP; DAKO, USA), according to manufacturer's instructions. Diaminobenzidine was used as chromogen.

**Immunohistochemical evaluation**

The expression of VGF was independently evaluated by one pathologist (F.S.) based on grading systems previously established for other markers (14, 15). Positive staining of VGF was considered when detected in the cytoplasm of cancer cells and in the tumor-adjacent stroma. The cases were classified as negative/low when scored as (0), (1+), (2+), whereas (3+) were determined as high expressing cases. Additionally, VGF expression in the stroma was classified as positive when present and negative when absent. VGF positive was considered when we found VGF high expression (3+) in cancer cells and positive in the tumor-adjacent stroma.

**Statistical analyses**

Error bars in graphical data represent mean ± standard error of the mean (SEM). Reference to the number of independent biological replicates performed for each experiment, as well as the sample size of each experimental group/condition, is provided in the respective figure legend of each experiment. Statistical significance was determined with one-way ANOVA, in which P values smaller than 0.05 were considered statistically significant. The variance was similar between the groups that were statistically compared. Graph Pad Prism version 9.0c software (Graph Pad Software, San Diego, CA, USA) was used for statistical analysis and graphical data presentation. ImageJ was used for image processing and analysis. Survival analyses for the primary breast tumor were estimated by the Kaplan–Meier method and compared using the log-rank test. For statistical analysis of the immunohistochemistry results, chi-square tests were carried out using IBM® SPSS® statistics V.26 software package for Windows (SPSS, Inc., Chicago, IL, USA). All statistical tests were two-sided.
